# EvoRMD: Integrating Biological Context and Evolutionary RNA Language Models for Interpretable Prediction of RNA Modifications

**DOI:** 10.64898/2026.03.22.713386

**Authors:** Bo Wang, Hao Zhang, Taoyong Cui, Xiaoyu Wang, Jiangning Song, Hao Xu

## Abstract

RNA modifications are essential regulators of post-transcriptional gene expression, influencing RNA stability, localization, translation, and degradation. Determining the specific modification at a given nucleotide is therefore critical for understanding its regulatory role. Most computational approaches treat each modification type as an independent binary task. This strategy provides a macro-level statistical perspective, but it does not reflect that, under a defined biochemical or cellular condition, only one modification type can occur at a specific site. Current mapping assays also report a single observed modification per site, leaving all other types unlabeled rather than truly negative. These properties motivate a framework that can reason over competing modification types. We introduce **EvoRMD**, a unified model for biologically contextualized and interpretable prediction of RNA modification types. EvoRMD integrates contextual sequence embeddings from a large-scale RNA language model with structured biological metadata—including species, organ, cell type, and subcellular localization. A lightweight attention mechanism highlights informative sequence positions. A shared multi-class classifier then generates a context-conditioned plausibility distribution over eleven modification types (Am, Cm, Um, Gm, D, pseudouridine, m^1^A, m^5^C, m^5^U, m^6^A, m^7^G), consistent with the single-positive, multiple-unlabeled nature of existing datasets. Although trained in a multi-class setting, EvoRMD also produces calibrated multi-label predictions through sigmoid-transformed logits, enabling direct comparison with existing single-modification and multi-label methods. EvoRMD achieves strong predictive performance and offers interpretable insights through attention profiles and motif analyses. Together, these components establish a biologically grounded framework for identifying and prioritizing RNA modification types from sequence and context.

## 1 Introduction

Since the first discovery of RNA modifications, more than 170 RNA modifications have been identified in the past six decades, collectively referred to as epitranscriptomes ^1,2^. These modifications are critical regulators of biological expression systems and represent some of the most conserved properties during evolution ^3^. Notably, modifications such as *N*^6^-methyladenosine (m^6^A), 5-methylcytosine (m^5^C), *N*^1^-methyladenosine (m^1^A), cytosine *N*^4^-acetylation (ac4C), dihydrouridine (D), *N*^7^-methylguanosine (m^7^G), 2’-O-methylation (Nm) and pseudouridine (Y) have been extensively characterized ^4,5^. These RNA modifications play important regulatory roles in various aspects of transcription and translation processes such as proper gene translation, RNA stability, ribosomal RNA translation, and protein translation efficiency and fidelity ^6,7,8,9^. Furthermore, dysregulation of RNA modifications has been increasingly linked to the onset and progression of diverse diseases, including cancer, neurological disorders, and metabolic syndromes ^10,11^. Therefore, accurate prediction of RNA modification types is essential for elucidating their functional roles in gene regulation, organismal development, and the pathogenesis of various diseases.

Computational methods have become indispensable tools for predicting RNA epitope modifications directly from primary sequences. Many existing models, such as SRAMP^12^, RNAm5CPred ^13^, m6AmPred ^14^, DeepBindGCN ^15^, Gene2vec ^16^, and DeepAc4C ^17^, have demonstrated high accuracy in identifying individual types of RNA modifications. However, current methodologies present fundamental limitations. First, most methods formulate each modification type as an independent binary classification task. This practice is driven not by biological considerations but by the modification-specific nature of early epitranscriptomic mapping assays, which detect only a single chemical mark per experiment and thus generate separate per-modification positive sets in resources such as RMBase and MODOMICS ^18,19,20,21^. Models built on such data implicitly assume that modification types occur independently and with similar priors. Yet extensive evidence shows that writer and eraser enzymes exhibit highly species-, tissue-, cell-type-, and subcellular-specific expression and localization patterns ^22,23,24,25,26,27,28^, and that several pathways are activated only under defined physiological or stress conditions (e.g., stress-induced pseudouridylation ^29^). RNA modifications therefore arise from context-dependent biochemical constraints rather than from independent Bernoulli processes, challenging the assumptions underlying binary formulations. Second, these approaches often rely on handcrafted sequence-derived features, such as k-nucleotide frequency (KNF), k-spaced nucleotide pair frequency (KSNPF), and pseudo-nucleotide composition (pseDNC) ^17,30,31,32,33,13^. While these features capture low-level sequence statistics, they insufficiently represent higher-order contextual, evolutionary, or structural signals, limiting predictive generalization. Third, while predictive performance metrics such as accuracy are frequently emphasized, the interpretability and biological credibility of the results are equally crucial for downstream applications. In particular, uncovering conserved sequence motifs associated with RNA modifications is essential for understanding the sequence-dependent regulatory mechanisms that govern epitranscriptomic modifications.

To address these challenges, we introduce **EvoRMD**, a unified framework for biologically contextualized and interpretable prediction of RNA modification types (Fig. 1). EvoRMD integrates RNA-FM sequence embeddings with structured biological context—including species, organ, cell type, and subcellular localization—to model the biochemical environments that influence modification specificity. A lightweight attention mechanism highlights informative sequence positions, and a shared classifier jointly evaluates all candidate modification types from a single representation. Current epitranscriptomic datasets provide a single observed modification per site, while all remaining types are unlabeled due to the modification-specific nature of existing mapping assays. Under this supervision structure, a multi-class formulation offers a principled and biologically coherent solution: it reflects the mutually exclusive nature of modification outcomes at a given nucleotide under specific biological conditions, avoids the artificial false-negative assumptions inherent in one-vs-rest binary models, and produces a context-conditioned plausibility distribution across all modification types. Although EvoRMD is optimized within this multi-class framework, the model outputs one logit per modification type. Through calibrated sigmoid transformation, EvoRMD also produces multi-label predictions, enabling direct comparison with existing single-modification and multi-label approaches. Collectively, these components establish a rigorous and biologically grounded framework for prioritizing RNA modification types from sequence and context.

**Fig. 1.**
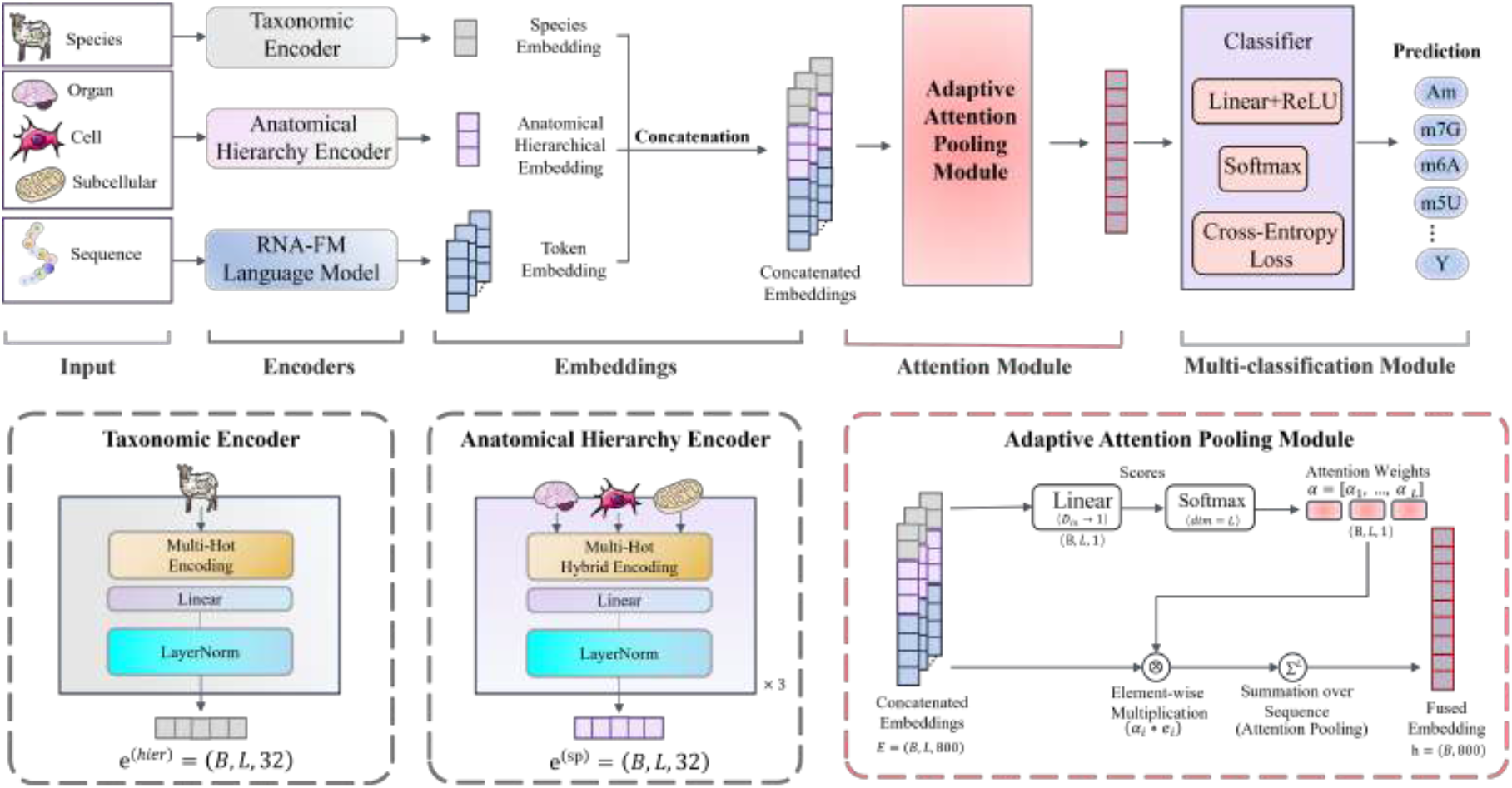
The framework of EvoRMD. The model consists of four main components: Input Sequences, Multi-branch Embedding Module, Attention Module, and Multi-classification Module. Given an RNA sequence centered on a candidate modification site, contextual embeddings are obtained from the RNA-FM language model, while complementary biological context features—including hierarchical (organ, cell, and subcellular) and species embeddings—are generated through parallel hybrid multi-hot encoders. These representations are concatenated and fused into a unified feature space (B,L,800). A trainable attention mechanism assigns position-wise weights to highlight informative regions and aggregates the sequence features into a global representation. The aggregated embedding is then passed to a multi-layer perceptron (MLP) classifier for joint prediction of multiple RNA modification types, enabling EvoRMD to integrate molecular sequence information with biological context for improved interpretability and prediction accuracy.

## 2 Results

### 2.1 Overview of EvoRMD framework

The EvoRMD framework integrates multi-scale biological context with RNA sequence information through a unified and interpretable architecture Fig. 1. The model is composed of an input module and three encoder branches—the Taxonomic Encoder, the Anatomical Hierarchy Encoder, and the RNA-FM Language Model—which are followed by a Adaptive Attention Pooling Module and a Unified Multi-Classification Module. Together, these components enable EvoRMD to seamlessly integrate evolutionary, anatomical, and sequence-derived information into a coherent representation for accurate RNA modification prediction.

The input to EvoRMD comprises a 41-nucleotide RNA sequence window (20 nt upstream and downstream) augmented with biological metadata—including species, organ, cell type, and subcellular localization—that provide multi-scale contextual signals known to influence RNA modification machinery. The Taxonomic Encoder converts species information into multi-hot vectors and projects them through a linear layer with normalization, generating a compact taxonomic embedding. The Anatomical Hierarchy Encoder processes organ, cell type, and subcellular location independently using hybrid multi-hot–hashing embeddings, producing three anatomical embeddings that capture hierarchical biological structure. The RNA sequence itself is encoded using RNA-FM ^34^, a pretrained 12-layer transformer language model. For each nucleotide position, RNA-FM outputs contextual embeddings that capture local and long-range sequence dependencies. These frozen representations serve as the primary sequence-based features. The trainable attention mechanism fuses taxonomic, anatomical, and RNA-FM embeddings into a unified token representation by assigning position-specific importance scores via a linear transformation and softmax normalization. The resulting attention weights guide a weighted aggregation of the fused embeddings, yielding a global sequence representation that emphasizes biologically meaningful positions and metadata-linked contextual signals. The aggregated representation is fed into a lightweight multi-layer perceptron classifier that outputs a probability distribution over eleven RNA modification types (Am, Cm, Um, Gm, D, Y, m^1^A, m^5^C, m^5^U, m^6^A, or m^7^G). By adopting a unified multi-class formulation rather than multiple binary classifiers, EvoRMD learns discriminative relationships between modification types and avoids redundancy introduced by independent classifiers. Overall, the architecture tightly integrates species-to-subcellular biological context, RNA structural dependencies captured by RNA-FM, and attention-based feature fusion, resulting in a robust and interpretable framework for multi-type RNA modification prediction.

### 2.2 EvoRMD performance

Before presenting the performance results, we briefly clarify the task setting (see Section 5.1 for details). Although multiple RNA modifications can in principle occur at the same nucleotide, the RMBase benchmark used in this study provides only a single observed modification label per site. This arises from the data-generating process: current mapping assays detect one modification type per experiment, leaving all remaining types *unobserved* rather than true negatives. Under this resulting single-positive, multiple-unlabeled supervision structure, a single-label multi-class formulation with softmax cross-entropy is the appropriate and loss-consistent choice for model training. We therefore first evaluate EvoRMD in this single-label multi-class setting and then show, within the same section, how the same trained model can also yield complementary multi-label predictions without retraining.

#### 2.2.1 Multi-class classification performance

EvoRMD demonstrates strong overall performance under the standard multi-class setting, achieving high accuracy and well-balanced precision–recall characteristics across most modification categories (Table 1). Widely occurring modifications with abundant training signals—such as m^6^A, m^5^C, Um, m^7^G, and Gm—reach MCC values above 0.95, confirming the model’s ability to extract highly discriminative sequence and contextual features. Even for moderate- and low-frequency categories such as m^1^A and D, EvoRMD maintains robust performance, achieving MCC values above 0.87. Nevertheless, the model exhibits varying levels of difficulty across different modification types. For example, Am displays comparatively lower precision and recall (79.17% and 73.08%, respectively), yielding a moderate MCC of 75.97%. This reduction likely arises from the subtle sequence signals that characterize Am and its overlap with nearby adenosine-centered modifications such as m^1^A and m^6^A. In contrast, rare modifications such as m^5^U—despite limited sample availability—still reach perfect accuracy or recall at the softmax stage, indicating that EvoRMD can capture highly discriminative local sequence or biological-context features sufficient for reliable identification of even scarce modification types.

**Table 1.** Per-class performance (averaged over three runs) for 11 RNA modifications.

| Type | Accuracy(%) | Precision(%) | Recall(%) | F1-score(%) | MCC(%) |
| --- | --- | --- | --- | --- | --- |
| Am | 73.08 | 79.17 | 73.08 | 76.00 | 75.97 |
| Cm | 94.52 | 94.52 | 94.52 | 94.52 | 94.46 |
| D | 90.91 | 83.33 | 90.91 | 86.96 | 87.02 |
| Gm | 100.00 | 94.59 | 100.00 | 97.22 | 97.24 |
| m<sup>1</sup>A | 93.42 | 91.03 | 93.42 | 92.21 | 92.12 |
| m<sup>5</sup>C | 98.96 | 98.96 | 98.96 | 98.96 | 98.90 |
| m<sup>5</sup>U | 100.00 | 50.00 | 100.00 | 66.67 | 70.69 |
| m<sup>6</sup>A | 99.88 | 99.88 | 99.88 | 99.88 | 99.08 |
| m<sup>7</sup>G | 97.62 | 100.00 | 97.62 | 98.80 | 98.79 |
| Um | 100.00 | 99.32 | 100.00 | 99.66 | 99.65 |
| Y | 86.11 | 100.00 | 86.11 | 92.54 | 92.76 |
| **Overall** | 99.47 | 99.49 | 99.47 | 99.47 | 97.82 |

To further understand these performance differences, we analyzed the softmax confidence distributions (Fig. 2 a). High-performing categories (Gm, m^6^A, m^5^C, Y, Um) exhibit narrow, right-skewed confidence distributions concentrated near 1.0, reflecting strong model certainty and clear separability. In contrast, Am displays a wider and more symmetric distribution with a lower median confidence, consistent with its reduced MCC and indicating intrinsic ambiguity in distinguishing Am from neighboring modification classes. The prediction flow visualized by the Sankey diagram (Fig. 2 b) further supports these observations. Most categories show tightly aligned links between true and predicted labels, demonstrating stable mapping in the multi-class setting. Misclassifications are largely concentrated in a small number of biologically related pairs, particularly Am vs. m^1^A or m^6^A and Cm vs. m^5^C, which correspond to known biochemical similarities and partially overlapping sequence contexts. These structured flow patterns indicate that the model’s errors follow systematic and biologically meaningful patterns rather than arising from random noise. EvoRMD demonstrates strong one-vs-rest discrimination across all 11 RNA modification types, as shown by the ROC and PR curves in Fig. 2 c-d. The ROC curves for all categories lie close to the upper-left boundary, indicating high true-positive rates accompanied by low false-positive rates, which reflects the model’s ability to distinguish each modification type from the remaining classes. The PR curves display a similarly favorable pattern, with most curves concentrated toward the upper-right region of the plot, demonstrating high precision at high recall. This behavior is particularly meaningful under strong class imbalance, where PR curves provide a more sensitive assessment of minority-class performance. Overall, the combined results from Table 1 and Fig. 2 indicate that EvoRMD, powered by a multi-source embedding architecture that integrates contextual representations from RNA-FM with taxonomic and anatomical hierarchy encodings, together with a lightweight attention mechanism and a unified MLP softmax classifier, achieves robust and biologically consistent multi-class performance.

**Fig. 2.**
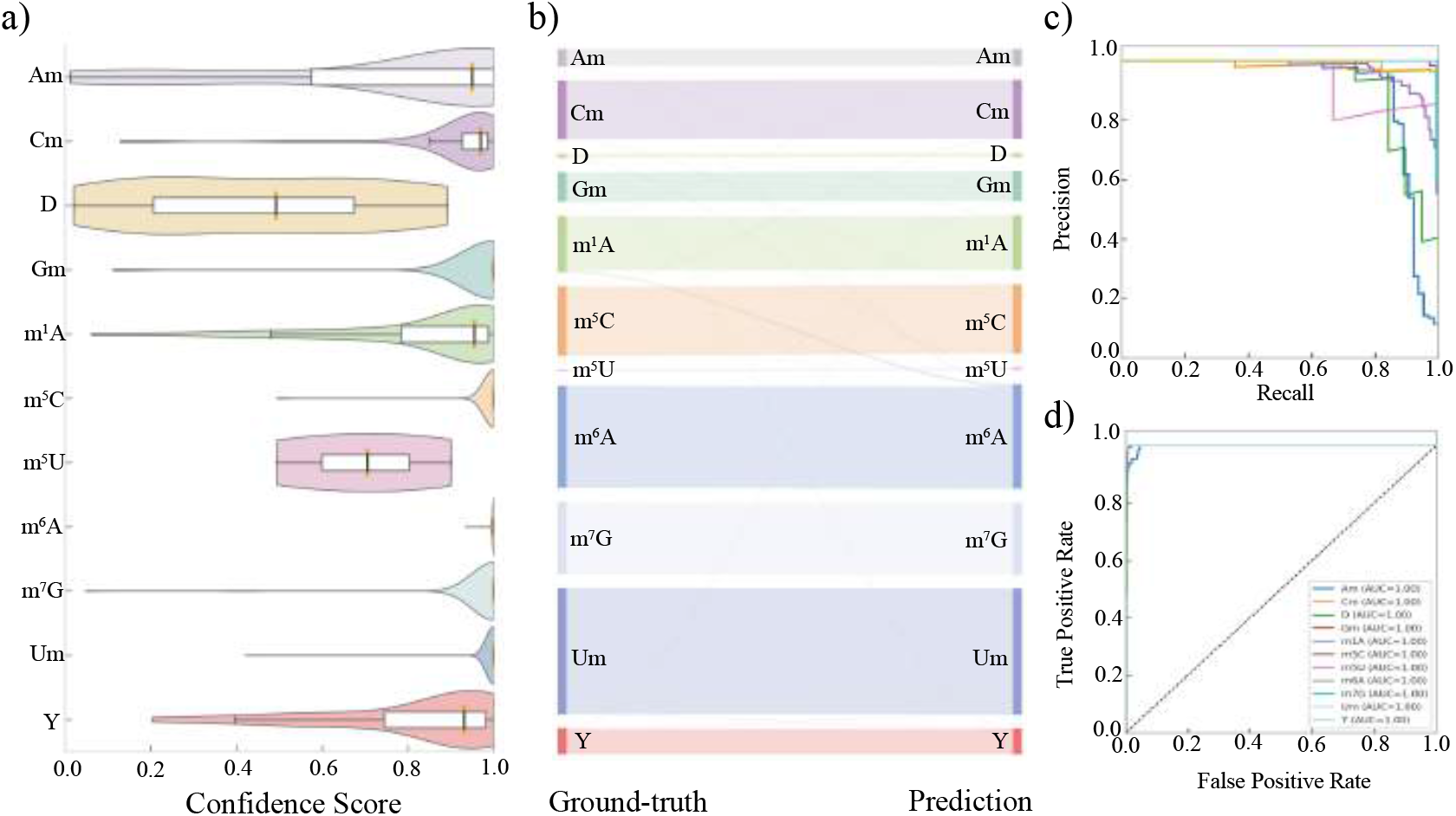
**a)** Confidence score distributions for all 11 RNA modification types generated by the softmax output of the multi-class classifier. Most categories display right-skewed, high-confidence distributions, whereas a few modifications show broader or lower-confidence patterns, reflecting varying levels of separability. **b)** Sankey diagram illustrating the mapping between ground-truth labels and predicted classes in the multi-class setting. Most flows align tightly, indicating stable classification behavior, while the few cross-category flows correspond to biologically related modification pairs. **c)** Precision–Recall (PR) curves for the one-vs-rest evaluation of each modification type. The curves show high precision at high recall for the majority of classes, demonstrating strong discriminative ability even under class imbalance. **d)** Receiver Operating Characteristic (ROC) curves for all modification types under the one-vs-rest setting. All classes achieve near-perfect ROC profiles with AUC values approaching 1.0, indicating excellent separability across categories. All results are produced using EvoRMD’s unified multi-class prediction framework with a softmax output over the 11 RNA modification types.

#### 2.2.2 Multi-label classification performance

In addition to the single-label multi-class evaluation, we also derive multi-label predictions from the same trained EvoRMD model to examine the compatibility patterns among modification types. This is done without retraining: the raw logits are transformed with sigmoid to yield independent likelihoods for each modification type, followed by per-class threshold calibration on the validation set using one-vs-rest MCC. The resulting modification-specific thresholds (Table 2), typically in a high-confidence range (*t*^∗^ ≈ 0.84–1.00), produce biologically interpretable multi-label outputs on the test set. As shown in Table 2, most classes achieve remarkably high MCC values after calibration, with eight modifications exceeding an MCC of 90%, and several—such as m6A, m5U, Um, and Cm—approaching near-perfect discriminability. These strong MCC values demonstrate that EvoRMD not only maintains accurate single-label predictions but also generalizes effectively in a multi-label setting, where modification types are evaluated independently.

**Table 2.** Optimal thresholds and MCC values for each modification in multi-label calibration.

| Modification | Am | Cm | D | Gm | m<sup>1</sup>A | m<sup>5</sup>C | m<sup>5</sup>U | m<sup>6</sup>A | m<sup>7</sup>G | Um | Y |
| --- | --- | --- | --- | --- | --- | --- | --- | --- | --- | --- | --- |
| **Optimal threshold** | 0.99 | 0.99 | 0.84 | 0.97 | 0.99 | 0.99 | 0.94 | 0.99 | 0.96 | 0.99 | 0.99 |
| **Best MCC (%)** | 86.79 | 97.96 | 91.67 | 95.89 | 94.34 | 92.55 | 100.00 | 99.76 | 96.00 | 99.32 | 95.77 |

Using these calibrated thresholds, we generated multi-label predictions for each RNA sequence and examined how often different modification types tend to be predicted together. The resulting co-occurrence bubble matrix (Fig. 3) shows a clear and intuitive pattern. Most of the signal is concentrated along the diagonal, meaning that EvoRMD continues to identify the true modification type with the highest confidence even after switching from a single-label softmax setting to an independent multi-label formulation. Beyond these main diagonal signals, we also observe several mild but meaningful off-diagonal associations. For example, Am occasionally appears together with m^1^A or m^6^A, which is expected because all three modifications occur at adenosines and share similar local sequence features. Likewise, Cm sometimes co-occurs with m^5^C, a pair known to be diffcult to distinguish in bisulfite-based detection assays ^35^. These associations do not occur randomly; instead, they appear only in a few biologically plausible pairs, indicating that EvoRMD is detecting subtle similarities between related modification types rather than generating widespread false positives. By extending EvoRMD from a softmax-based multi-class setting to a multi-label framework, we gain an additional perspective on what the model has learned. After converting the logits into independent probabilities using sigmoid and applying class-specific thresholds, we can see which modification types tend to appear together and how strongly the model supports each label. This analysis reveals signals that are not visible under single-label softmax training alone. Overall, extending EvoRMD into a multi-label prediction framework reveals an additional layer of interpretability that complements the strong softmax classification performance. The calibrated sigmoid outputs expose latent modification-specific compatibility signals encoded in the logits, enabling the model to reflect biological relatedness among modification types and supporting downstream tasks such as motif comparison, modification neighborhood analysis, or condition-specific modification co-regulation studies. Together, the softmax and multi-label evaluations demonstrate that EvoRMD not only predicts RNA modification types with high accuracy but also learns a coherent biochemical landscape embedded within the modification space.

**Fig. 3.**
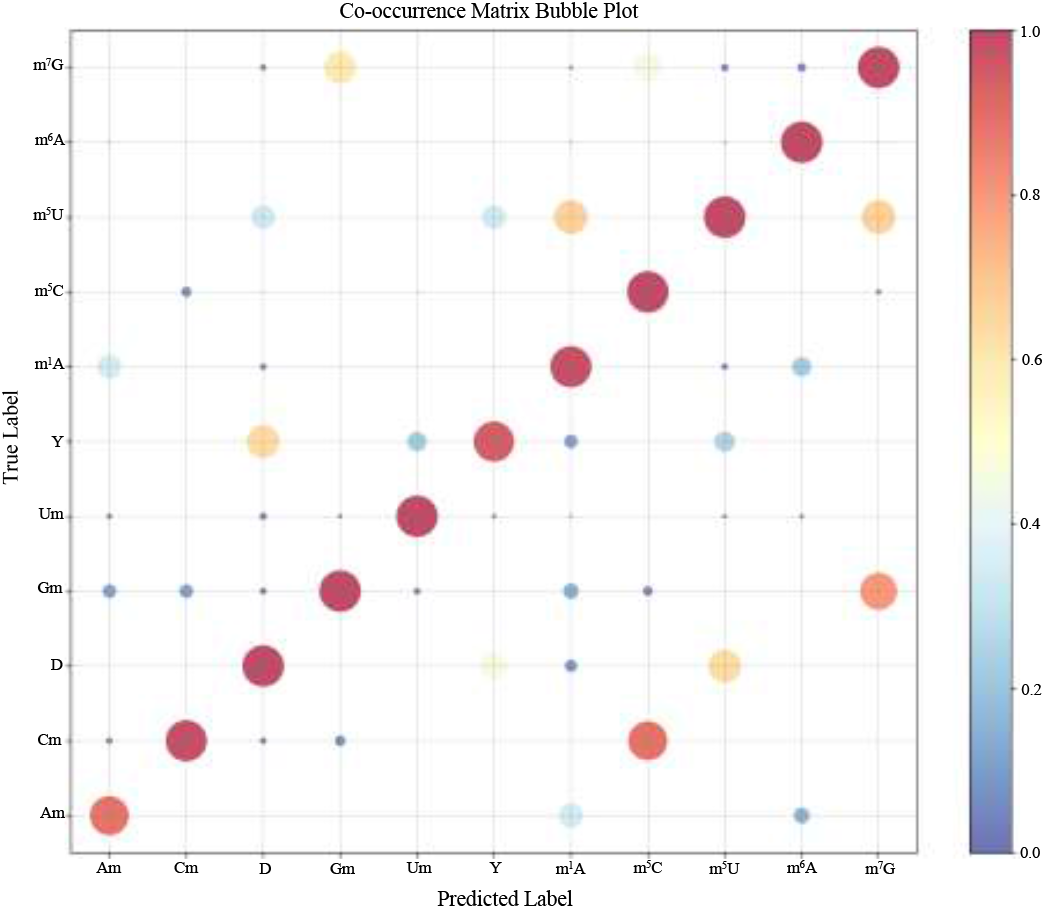
Co-occurrence bubble plot of multi-label predictions generated by EvoRMD. Each bubble represents the fraction of samples belonging to a true modification type (rows) that are predicted as positive for a given modification label (columns) after sigmoid calibration and class-specific thresholding. Bubble size indicates the co-occurrence frequency, and color reflects the normalized value (0–1). Strong diagonal signals indicate that EvoRMD consistently assigns the correct modification type in the multi-label setting. A few mild off-diagonal signals appear only in biologically related modification pairs—for example, Am with m^1^A/m^6^A and Cm with m^5^C—reflecting shared sequence features or known limitations of experimental detection methods. Overall, the matrix illustrates that EvoRMD captures both accurate primary predictions and meaningful secondary compatibility patterns among RNA modification types.

### 2.3 Comparison with state-of-the-art methods

EvoRMD supports two evaluation modes—multi-class and multi-label—and we compare it with baselines appropriate for each. In the multi-class setting, EvoRMD is benchmarked against multi-modification models; in the multi-label setting, it is compared with state-of-the-art single-modification predictors.

#### 2.3.1 Multi-class setting: comparison with multi-modification models

We first compare EvoRMD with two state-of-the-art multi-modification predictors, MultiRM ^36^ and TransRNAm ^37^, which represent the strongest existing frameworks capable of jointly predicting ten or more RNA modification types. Both baselines adopt a multi-class architecture composed of independent sigmoid-based classifiers—one for each modification type—and report performance under a per-class one-vs-rest AUROC protocol. For fair comparison, we follow exactly the same evaluation strategy: for each modification type, all positive sites are contrasted against all remaining sites, and a threshold-free AUROC is computed. EvoRMD’s softmax probability for the corresponding class is used directly as the ranking score in the same one-vs-rest analysis, ensuring that architectural differences between sigmoid-based and softmax-based formulations do not affect the fairness of the comparison. As shown in Table 3, EvoRMD achieves the highest AUROC for all 11 RNA modification types, yielding an average AUROC of 0.9940 compared with 0.9432 for MultiRM and 0.9572 for TransRNAm. Figure 4 further quantifies these improvements: relative AUROC gains range from 0.69% to 11.18% over MultiRM and from 0.69% to 6.12% over TransRNAm, with particularly notable improvements for m^1^A, Y, Um, and m^5^U. Paired *t*-tests confirm that these improvements are statistically significant (EvoRMD vs. MultiRM: *p* = 0.0012; EvoRMD vs. TransRNAm: *p* = 0.0005). Importantly, EvoRMD outperforms both baselines across all modification types, indicating a broad and systematic advantage rather than isolated gains on a few categories.

**Table 3.** AUC comparison of RNA modifications across EvoRMD, MultiRM, and TransR-NAm models.

| Modification | EvoRMD | MultiRM | TransRNAm |
| --- | --- | --- | --- |
| Am | **0.9982** | 0.8954 ($\downarrow 0.1028$) | 0.9402 ($\downarrow 0.0580$) |
| Cm | **0.9999** | 0.9663 ($\downarrow 0.0336$) | 0.9692 ($\downarrow 0.0307$) |
| Gm | **1.0000** | 0.9805 ($\downarrow 0.0195$) | 0.9847 ($\downarrow 0.0153$) |
| Um | **1.0000** | 0.9440 ($\downarrow 0.0560$) | 0.9511 ($\downarrow 0.0489$) |
| D | **0.9998** | — | — |
| Y | **0.9999** | 0.9372 ($\downarrow 0.0627$) | 0.9386 ($\downarrow 0.0613$) |
| m<sup>1</sup>A | **0.9996** | 0.8975 ($\downarrow 0.1021$) | 0.9566 ($\downarrow 0.0430$) |
| m<sup>5</sup>C | **1.0000** | 0.9679 ($\downarrow 0.0321$) | 0.9747 ($\downarrow 0.0253$) |
| m<sup>5</sup>U | **1.0000** | 0.9462 ($\downarrow 0.0538$) | 0.9556 ($\downarrow 0.0444$) |
| m<sup>6</sup>A | **0.9999** | 0.9875 ($\downarrow 0.0124$) | 0.9930 ($\downarrow 0.0069$) |
| m<sup>7</sup>G | **0.9999** | 0.9644 ($\downarrow 0.0355$) | 0.9666 ($\downarrow 0.0333$) |
| **Avg.** | **0.9940** | 0.9432 ($\downarrow 0.0508$) | 0.9572 ($\downarrow 0.0368$) |

Note: The best result is highlighted in **bold**. Green arrows indicate improvement over EvoRMD; red arrows indicate lower performance compared with EvoRMD.

**Fig. 4.**
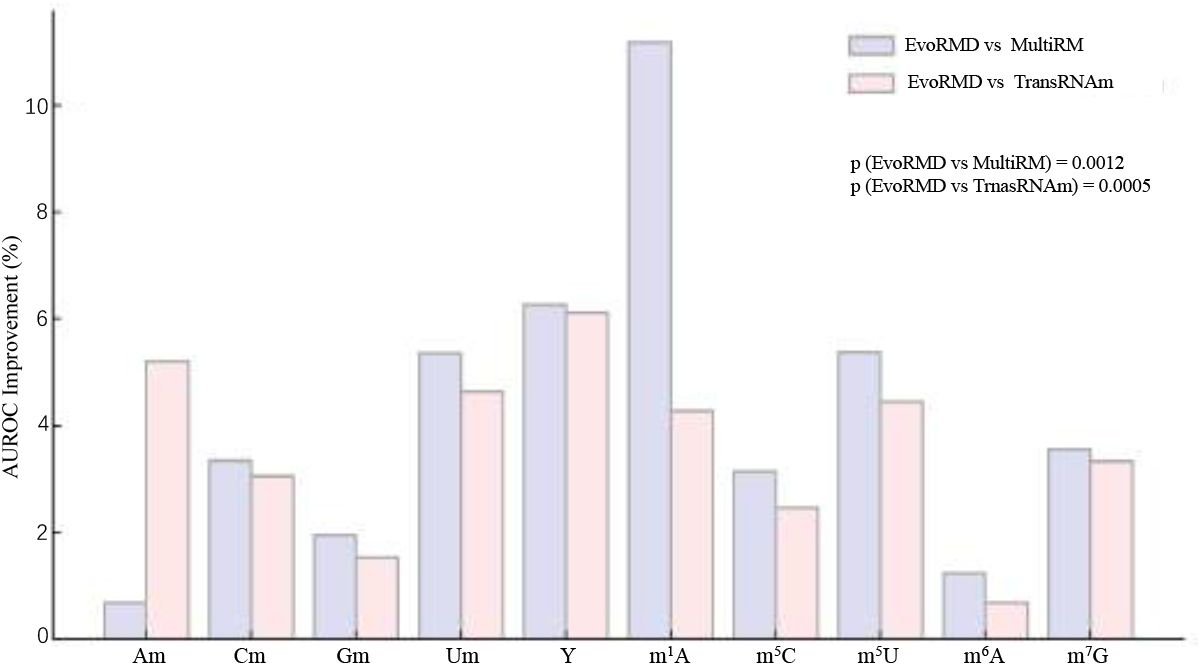
Relative AUROC improvement of EvoRMD over MultiRM and TransRNAm for 11 RNA modifications. AUROC values of EvoRMD are averaged over three seeds (42, 215, 3407). EvoRMD achieves consistent gains across all modifications, with mean improvements of 5.08% (vs. MultiRM) and 3.68% (vs. TransRNAm). Improvements are statistically significant (MultiRM: *p* = 0.0012, TransRNAm: *p* = 0.0005).

#### 2.3.2 Multi-label setting: comparison with single-modification models

In addition to the unified multi-class evaluation, we also assess EvoRMD in a multi-label setting, where each modification type is treated as an independent prediction task. This setting enables direct comparison with existing single-modification predictors, many of which are designed specifically for one RNA modification and report performance only in a binary (positive vs. negative) classification framework. These task-specific models—including 2OMe-LM for Nm, m1A-Ensem for m^1^A, m5C-iEnsem for m^5^C, Stack-DHUpred for D, m5U-HybridNet for m^5^U, m6A-TSHub for m^6^A, MCAMEF-BERT for m^7^G, and Definer for Y the strongest available baselines for single-modification prediction (Table 4). To obtain multi-label predictions from EvoRMD, we apply a sigmoid transformation to the logits of the trained model and determine modification-specific thresholds on the validation set. This yields eleven independent 0/1 decisions, allowing EvoRMD to function equivalently to eleven parallel binary classifiers—one for each RNA modification type. Under this unified protocol, EvoRMD can be directly benchmarked against single-task models using the same MCC metric. As shown in Table 4, EvoRMD achieves competitive or superior performance across nearly all modifications, substantially outperforming published predictors for Nm, D, m^7^G, and Y, and performing comparably to the ensemble-based m1A-Ensem on m^1^A. These results demonstrate that EvoRMD, despite being trained as a single unified model, matches or exceeds highly specialized single-modification predictors when evaluated in a multi-label framework.

**Table 4.** Performance comparison between EvoRMD and state-of-the-art single-modification predictors under an independent multi-label evaluation.

| Modification | Best Single-Task Model | Reported MCC | EvoRMD | ΔMCC |
| --- | --- | --- | --- | --- |
| Nm (Am/Cm/Gm/Um) | 2OMe-LM<sup>38</sup> | 0.695 | 0.950* | +0.255 |
| m<sup>1</sup>A | m1A-Ensem<sup>39</sup> | 0.980 | 0.943 | -0.037 |
| m<sup>5</sup>C | m5C-iEnsem<sup>40</sup> | 0.878 | 0.926 | +0.048 |
| D (DHU) | Stack-DHUpred<sup>41</sup> | 0.567 | 0.917 | +0.350 |
| m<sup>5</sup>U | m5U-HybridNet<sup>42</sup> | 0.851 | 1.000 | +0.149 |
| m<sup>6</sup>A | m6A-TSHub<sup>43</sup> | 0.490 | 0.998 | +0.508 |
| m<sup>7</sup>G | MCAMEF-BERT<sup>44</sup> | 0.795 | 0.960 | +0.165 |
| Y | Definer<sup>45</sup> | 0.733 | 0.958 | +0.225 |

Note: \* The Nm performance of EvoRMD is the arithmetic mean of Am, Cm, Gm, and Um subtypes (MCC = 0.8546, 0.9776, 0.9556, and 0.9925, respectively). EvoRMD MCC values are computed from sigmoid-activated logits, and each modification type is evaluated under a 1:10 positive-to-negative sampling ratio.

### 2.4 Interpretation

RNA modification prediction is inherently multi-faceted, involving positional sequence preferences, conserved motifs, and potential relationships across modification types. In the following sections, we dissect the internal representations learned by EvoRMD in order to understand how the model captures biologically meaningful signals and whether these signals agree with known epitranscriptomic mechanisms.

#### 2.4.1 Analysis of attention weights learned by attention module

To identify which regions of the 41-nt window contribute most to EvoRMD’s predictions, we analyzed the attention weights from all true positive samples (Fig. 5). Because each nucleotide position is encoded by a fused representation combining RNA-FM sequence features with broadcasted biological metadata, the learned attention highlights where this integrated embedding is most informative for each modification type. As summarized in Table A1, the top-ranked positions are concentrated around the central site—most prominently position 20 and, for several modifications, position 22—together with a small set of recurrent upstream positions such as 3, 7, 15, and 19. The central nucleotide (N21) itself generally receives moderate but non-negligible attention; its average weight is systematically higher for A-, C-, and G-derived modifications (Am, Cm, Gm, m^1^A, m^5^C, m^6^A, m^7^G; N21 attention ≈ 0.021–0.024) than for U-based modifications (D, m^5^U, Um, Y; N21 attention ≈ 0.011–0.018), where the top-8 positions are dominated by flanking and upstream sites (e.g., 7, 15, 20, 24, 27, 29). This pattern is consistent with the heatmaps in Fig. 5, where A/C/G-related modifications show relatively smooth, symmetric peaks around the central region, whereas U-centered modifications exhibit sharper or shifted peaks toward neighboring nucleotides. Together, these observations suggest that EvoRMD relies more heavily on flanking-sequence cues to recognize U-centered modifications, while the modified nucleotide itself and its immediate neighborhood contribute more strongly to distinguishing A-, C-, and G-related modifications.

**Fig. 5.**
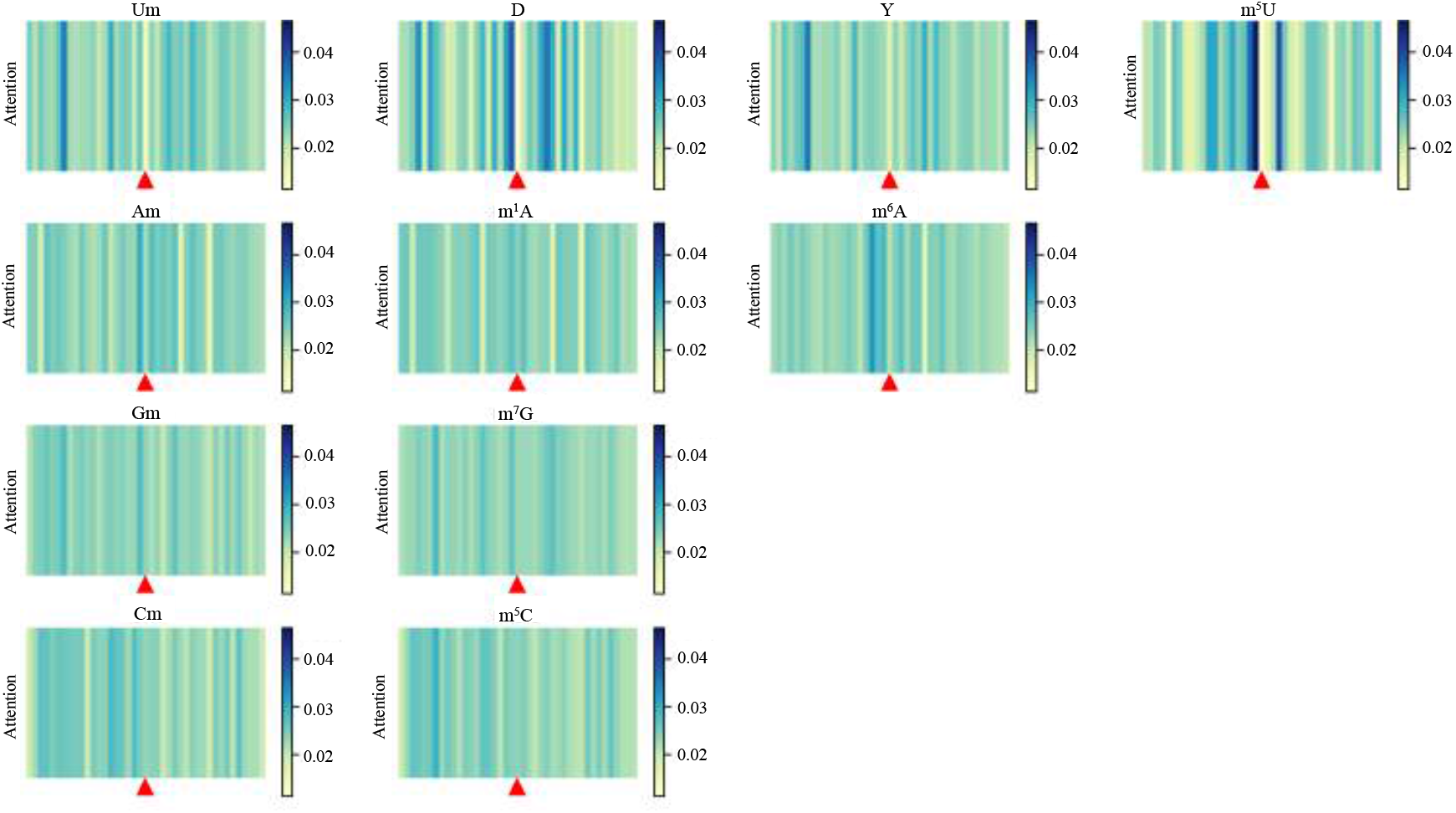
Attentional weighting patterns for different RNA modifications. It is obtained by averaging the weights of all TP samples of the same modification on the query axis. The x-axis represents the sequence direction, and the red triangle (▴) indicates the central (21st) nucleotide position that is modified.

Functional-region analysis further supports these patterns (Fig. 6). Modifications enriched in CDS, exons, and UTRs—such as Um, Y, Cm, and m^6^A—tend to exhibit dispersed attention distributions, aligning with their roles in translation efficiency and RNA stability that often rely on broader sequence context^46,47,48,49,50,51^. In contrast, modifications predominantly found in intronic or UTR regions—such as m^1^A and m^5^C—show more centralized, radially decaying attention around the modified site, consistent with localized sequence requirements for translation initiation, RNA stability control, and protein–RNA interactions ^52,47^. Taken together, these positional, global, and region-based analyses indicate that EvoRMD captures distinct yet biologically coherent recognition mechanisms across RNA modifications, with attention patterns that closely mirror known biochemical and functional behaviors.

**Fig. 6.**
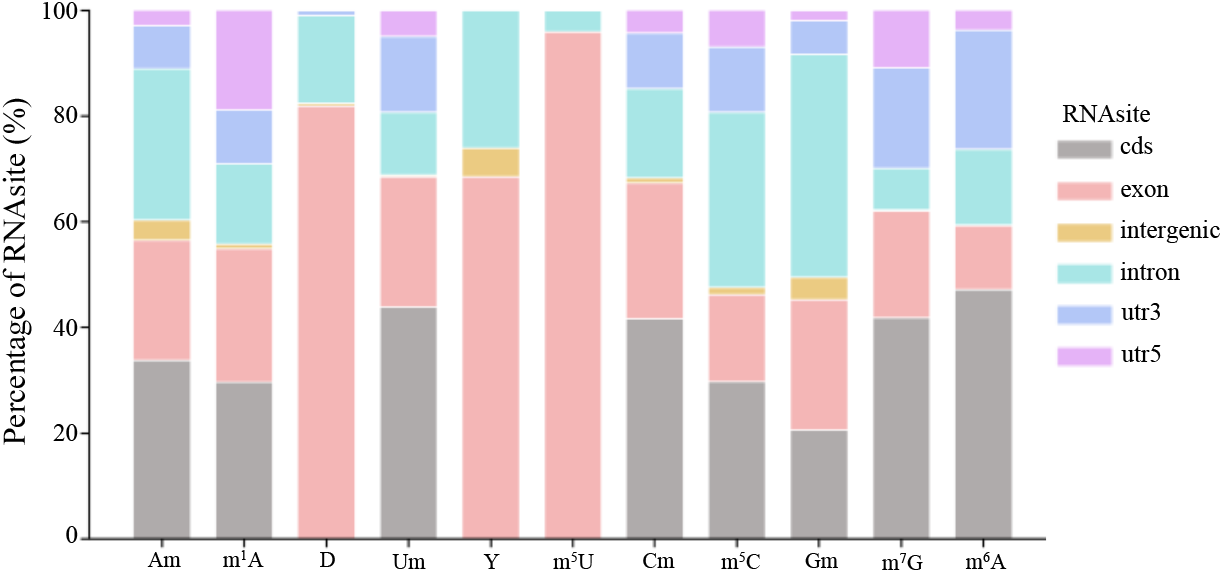
Genomic distribution of RNA modification sites across functional regions. Stacked bar plots show the proportion of each modification type located in coding sequence (CDS), exon, intron, 5’UTR and 3’UTR regions. These RNAsite annotations are used solely for post-hoc interpretation and are not provided as input features to EvoRMD. Modifications enriched in CDS, exonic and UTR regions—most notably Um, Y, Cm and m^6^A—tend to exhibit widely dispersed attention patterns, whereas those predominantly located in introns and UTRs—such as m^1^A and m^5^C—show more centralized, radially decaying attention around the modification site (Fig. 5). Together, these region-specific distributions provide an independent functional context for interpreting the attention patterns learned by EvoRMD.

#### 2.4.2 Conservative motif analysis

RNA modifications often exhibit conserved sequence motifs that have been maintained across evolution, reflecting their functional importance in guiding writer, reader, and eraser activities ^53^. To determine whether EvoRMD captures these underlying sequence regularities, we extracted high-attention windows for each modification type and transformed them into position-specific weight matrices (PWMs) representing putative motifs. As shown in Fig. 7, the motifs inferred by EvoRMD display clear correspondence to experimentally curated motifs in RMBase ^18^, with Tomtom ^54^ reporting statistically significant matches (*p<* 0.05) for the majority of modification types. All comparison results can be found in the appendix. Collectively, these results indicate that EvoRMD does not merely highlight salient nucleotides through attention, but captures evolutionarily conserved and functionally meaningful sequence patterns associated with RNA modification deposition, thereby reinforcing the biological validity and interpretability of the model.

**Fig. 7.**
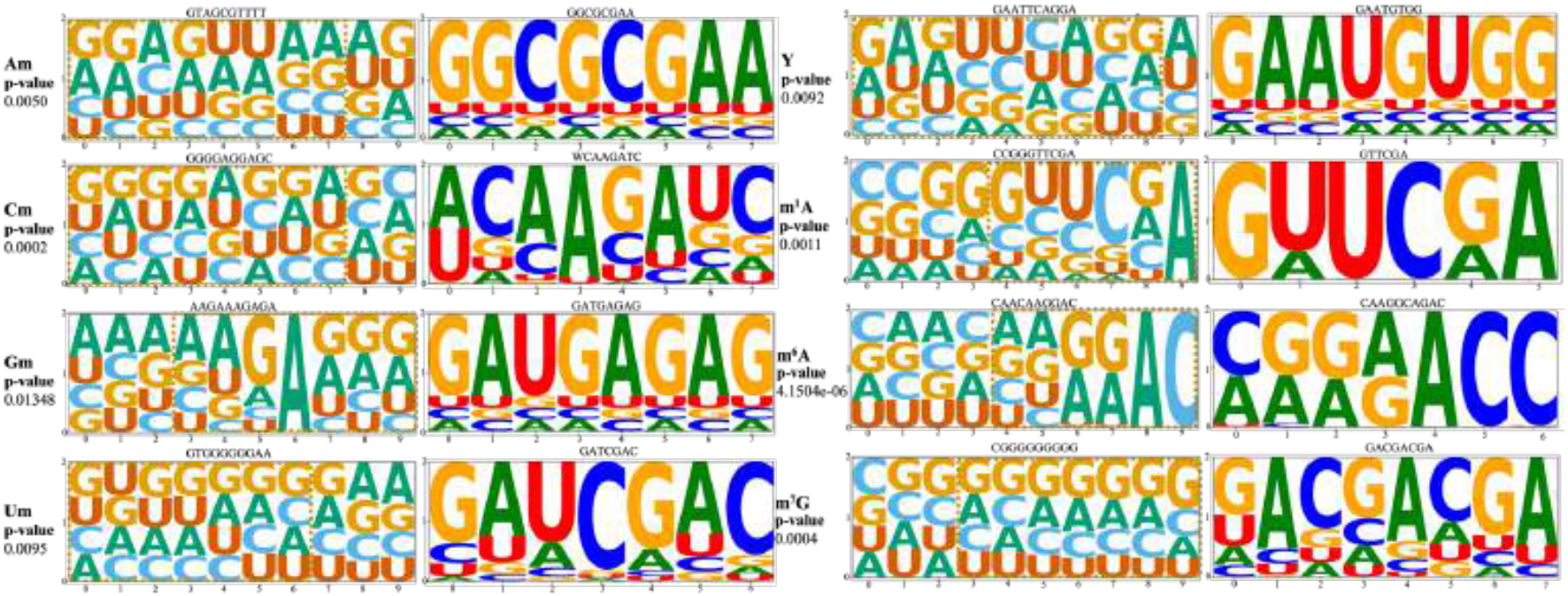
Comparison of EvoRMD-predicted motifs with known conserved RNA modification motifs from RMBase. Each motif pair consists of an EvoRMD-extracted sequence motif (left) and the most similar RMBase motif (right), as identified by Tomtom motif comparison. Motif similarity was quantified using Tomtom with a significance threshold of p *<* 0.05. The gold dashed boxes highlight the aligned regions of statistically significant similarity between the predicted and known motifs, suggesting that EvoRMD is capable of learning biologically meaningful and evolutionarily conserved sequence patterns associated with RNA modifications.

#### 2.4.3 Identify correlations between different modifications

Previous studies suggest that different RNA modifications may exhibit functional crosstalk ^55^. To assess whether EvoRMD captures such higher-order relationships, we first examined cross-modification motif similarity using both EvoRMD-derived motifs and curated motifs from RMBase ^18^. Tomtom ^54^ revealed widespread statistically significant matches (*p<* 0.05) across distinct modification types (Appendix B), indicating that several modifications share conserved sequence determinants. At the representation level, aggregation of RNA-FM ^34^ contextual embeddings around the modification site produced modification-specific vectors whose pairwise Pearson correlations showed strong positive relationships for most modification pairs (Fig. 8). Notably, high correlations were observed even between chemically distinct marks—such as m^5^U–Gm and m^6^A–m^7^G—suggesting potential shared biochemical or structural features encoded within the model’s learned representations.

**Fig. 8.**
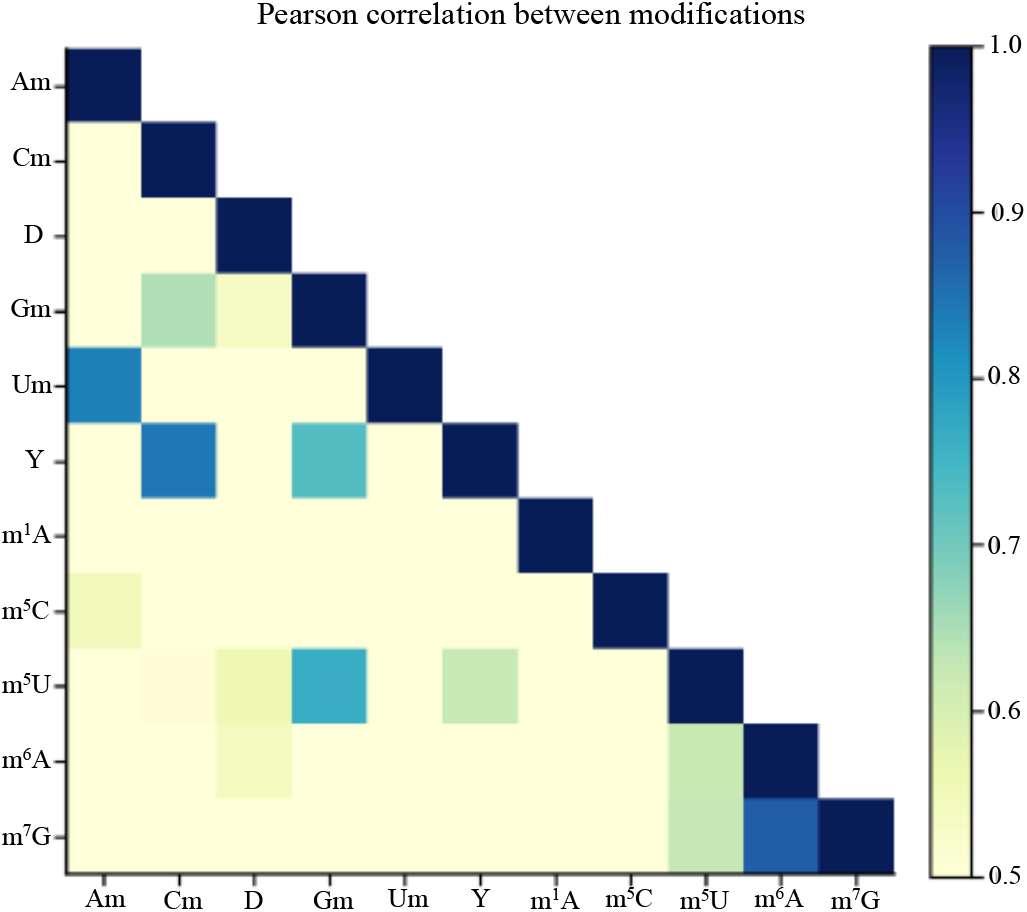
EvoRMD reveals associations between RNA modifications. The figure demonstrates pearson correlation among the weights of different RNA modificationsin the Attention module. Several pairs, such as m^7^G–m^6^A and Gm–m^5^U, show strong associations, indicating potential biological crosstalk or shared recognition mechanisms. These findings suggest that EvoRMD captures regulatory relationships beyond individual modifications.

To determine whether these relationships influence prediction behavior, we examined EvoRMD’s calibrated multi-label outputs. Although the model typically assigns a single dominant label, several modification pairs exhibited low-frequency but non-random co-occurrence patterns (Fig. 3), mirroring the embedding-level correlations. Examples include co-predictions of Am with m^1^A or m^6^A, and occasional Cm–m^5^C co-occurrence—pairs that are biochemically plausible due to shared nucleobase chemistry or recognition by overlapping writer–reader protein modules. These convergent patterns across motifs, embeddings, and predictions suggest that EvoRMD captures biologically coherent relationships that may arise from co-modification of proximal sites ^56^, modification-dependent structural priming ^55^, or reuse of the same enzymatic machinery across multiple marks ^57,58,59^.

#### 2.4.4 Dissecting the Contribution of Biological Context Components

To rigorously assess how multi-scale biological context—comprising species, organ, cell type, and sub-cellular localization—informs the model’s predictive capability, we conducted a systematic component analysis comparing the full EvoRMD framework against variants utilizing only specific context branches alongside the RNA-FM backbone. This decomposition allows us to disentangle the distinct roles played by evolutionary lineage information (Taxonomic Encoder) versus microenvironmental cues (Anatomical Hierarchy Encoder) in defining the RNA modification landscape.

We first visualized the latent feature spaces of these variants using t-SNE dimensionality reduction (Fig. 9). As observed in Fig. 9 a), the model integrating only species information (“Taxonomic + RNA-FM”) exhibits a feature topology similar to the sequence-only baseline (Fig. 9 c)), characterized by loose clustering and considerable overlap among modification types. This suggests that while evolutionary information provides a broad phylogenetic baseline, it is insufficient on its own to resolve the precise identity of modifications that are conserved across species but biochemically distinct. In strong contrast, the introduction of anatomical hierarchy information (“Anatomical + RNA-FM”, Fig. 9 b)) results in a drastic reorganization of the feature space, producing significantly tighter clusters and clearer decision boundaries. The full EvoRMD model (Fig. 9 d)) further refines this separation, yielding the most distinct topology. This visual evidence highlights that the fine-grained anatomical context acts as a primary driver for resolving modification ambiguities that sequence and species data alone cannot distinguish.

**Fig. 9.**
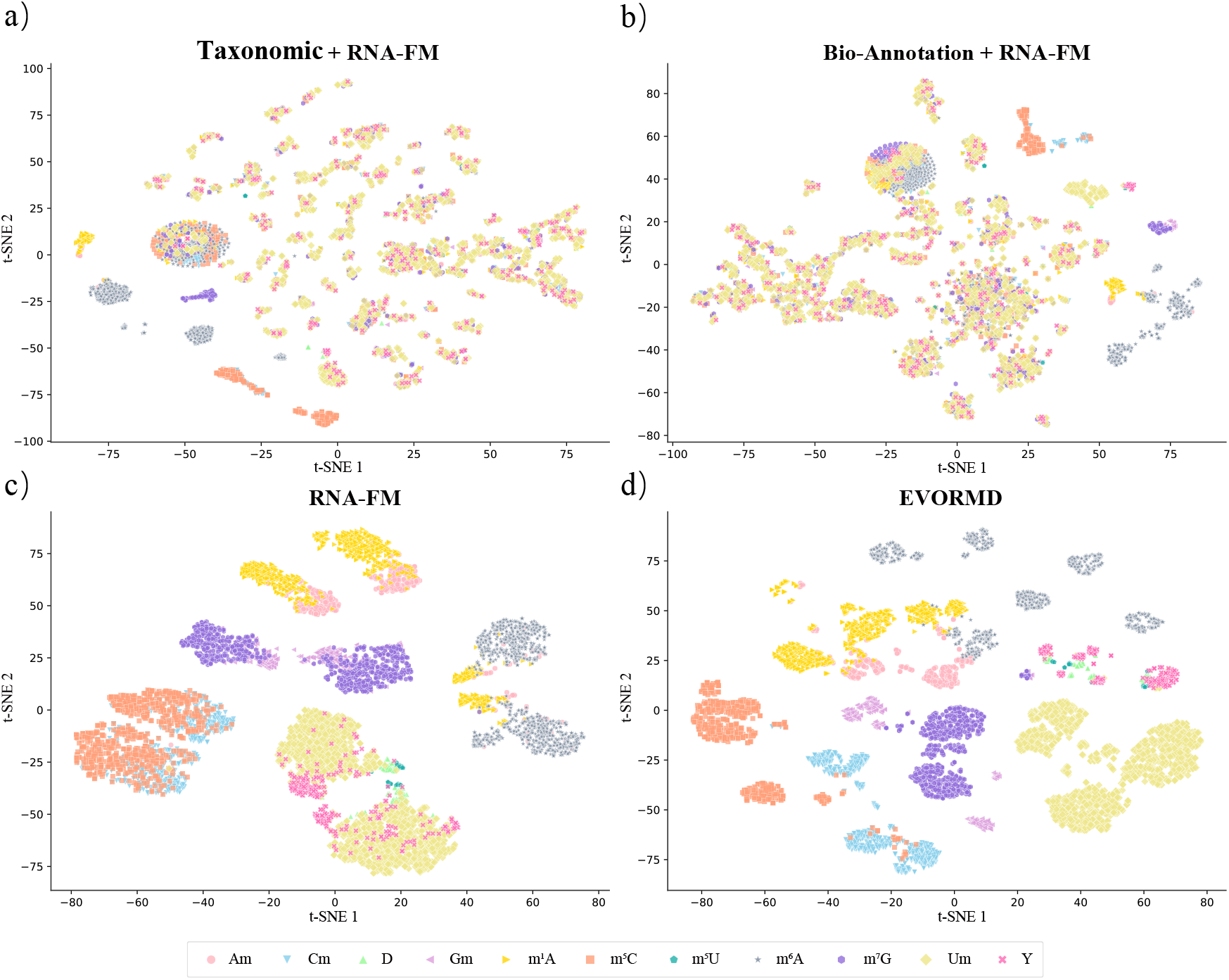
Visualization of the learned feature space using t-SNE dimensionality reduction. The figure compares the discriminative power of feature representations learned by four model variants on the test set. **a) Taxonomic + RNA-FM:** Feature space derived from integrating species information with RNA FM sequence embeddings. The clustering remains relatively loose with significant overlap, similar to the sequence-only baseline, indicating that species information alone provides limited discriminative gain. **b) Anatomical + RNA-FM:** Feature space generated by adding the Anatomical Hierarchy Encoder (organ, cell, subcellular) to RNA-FM. This variant shows drastically improved cluster separation compared to (a) and the baseline, highlighting the critical role of anatomical context in resolving modification ambiguities. **c) RNA-FM Only:** Baseline feature space using only pre-trained sequence embeddings, exhibiting fuzzy boundaries and overlap among biochemically similar modifications. **d) Full EvoRMD:** The final fused representation integrating Sequence, Taxonomic, and Anatomical features. This configuration yields the distinctest decision boundaries and tightest intra-class clustering, confirming that the comprehensive integration of evolutionary and anatomical signals maximizes discriminative power.

This observation is quantitatively substantiated by the classification performance (Table 5). The “Taxonomic + RNA-FM” variant yielded an overall MCC of 90.24%, which is comparable to the RNA-FM baseline (91.19%), reinforcing that species labels alone do not provide sufficient discriminative gain for site-specific prediction. Conversely, the “Anatomical + RNA-FM” variant achieved a notably higher overall MCC of 93.46%. Crucially, the anatomical encoder proved indispensable for modifications known to be highly regulated by tissue-specific enzyme expression or subcellular compartmentalization. For instance, in the prediction of Cm and Gm, the anatomical model achieved MCCs of 83.42% and 88.95%, respectively— far outperforming the taxonomic model (39.00% and 54.93%). Similarly, for Y, the anatomical context boosted the MCC to 94.59% compared to 69.31% with species info alone. However, the optimal performance was consistently achieved by the full EvoRMD framework (Overall MCC 97.82%), which integrates both encoders. This synergistic effect is most evident for m^1^A, where single-branch models struggled (MCC ∼44%), but the fused model achieved high accuracy (∼92%). These results demonstrate that RNA modifications are defined by a complex interplay of factors: while the Anatomical Hierarchy Encoder provides the critical resolution for tissue- and compartment-specific regulation, the Taxonomic Encoder anchors these signals within an evolutionary context. The comprehensive integration of both dimensions is therefore essential for the state-of-the-art performance and biological interpretability of EvoRMD.

**Table 5.** Quantitative assessment of the contribution of Taxonomic and Anatomical encoders. The table compares the Matthews Correlation Coefficient (MCC %) across four model configurations: the sequence-only baseline (RNA-FM), models with a single context branch (Taxonomic or Anatomical), and the complete EvoRMD framework.

| Modification | RNA-FM Only | Taxonomic + RNA-FM | Anatomical + RNA-FM | Full EvoRMD |
| --- | --- | --- | --- | --- |
| Am | **76.41** | 42.28 | 74.34 | 75.97 |
| Cm | 35.58 | 39.00 | 83.42 | **94.46** |
| D | 63.58 | 81.65 | **91.29** | 87.02 |
| Gm | 64.12 | 54.93 | 88.95 | **97.24** |
| $m^1A$ | 86.87 | 44.59 | 44.10 | **92.12** |
| $m^5C$ | 90.44 | 96.75 | **99.14** | 98.90 |
| $m^5U^*$ | **100.00** | 70.71 | 86.60 | 70.69 |
| $m^6A$ | 98.88 | 93.55 | 94.33 | **99.08** |
| $m^7G$ | 88.67 | 86.35 | 96.11 | **98.79** |
| Um | 93.23 | 94.26 | **99.68** | 99.65 |
| Y | 71.78 | 69.31 | **94.59** | 92.76 |
| **Overall MCC** | 91.19 | 90.24 | 93.46 | **97.82** |

Note: “Taxonomic + RNA-FM” integrates only species embeddings with the sequence backbone; “Anatomical + RNA-FM” integrates only hierarchical anatomical embeddings (organ, cell, subcellular). Bold numbers indicate the best performance. The variation in rare classes (e.g., $m^5U$ ) is influenced by sample size sensitivity.

### 2.5 Downstream profiling of motif patterns across cell types

To evaluate EvoRMD’s capacity to resolve epitranscriptomic context dependency, we performed a systematic downstream analysis focusing on the divergence and conservation of sequence motifs for modifications that are shared across cell lines with differing phenotypic and developmental backgrounds. We investigated the regulatory context of these shared RNA modifications by comparing motifs in two distinct cell line pairs: (1) two cancer cell lines of similar origin but divergent phenotypes (HepG2 and Huh7, derived from hepatocellular carcinoma), and (2) a pair representing a developmental transition from normal to cancerous states (human neural progenitor cells, HNPCs, and glioma stem cells, GSCs). This structured approach allowed us to differentiate between sequence features that are highly conserved due to essential enzymatic recognition and those that evolve to become cell-type–specific, reflecting the context-dependent nature of RNA modification regulation.

We selected these two pairs of cell lines to systematically evaluate EvoRMD’s ability to resolve cell-type-specific contextual features. The first pair, HepG2 and Huh7, are human hepatocellular carcinoma-derived epithelial cell lines of common tissue origin but divergent phenotypes, and share both m^1^A and m^6^A modification sites. The second pair, HNPCs and GSCs, represents a transition from normal to cancerous neural states and shares only the m^6^A modification. Despite their partial similarities, these cell lines exhibit well-documented differences in differentiation state, lipid and xenobiotic metabolism, viral susceptibility (e.g., HCV), and heterogeneous expression gradients of RNA modification writers, erasers, and readers ^60,61,62,63,64^. Multiple studies profiling m^6^A writers, erasers and readers across hepatocellular carcinoma cell lines (including HepG2 and Huh7) report heterogeneous expression and regulation of these factors, suggesting line-specific epitranscriptomic landscapes rather than a uniform pattern ^65,66,67,68,69^. Such backgrounds provide an ideal test case to examine whether EvoRMD learns cell-type–specific contextual features within the same RNA modification type. Common to all pairs, we utilized EvoRMD’s attention mechanism to analyze the shared modification sites. For each cell line and modification type, the top-10 attention-enriched local sequence windows were extracted, and independent motif discovery was performed. A cross-cell line comparison using Tomtom ^54^ on these extracted motifs revealed distinct patterns of evolutionary pressure and cell-type specificity.

#### 2.5.1 Intra-Cancer Heterogeneity (HepG2 vs Huh7)

The analysis of the shared m^1^A and m^6^A modifications between HepG2 and Huh7 revealed a clear difference in evolutionary pressure and cell-type specificity between the two modifications (Figure 10 a). For m^1^A, the attention-enriched motifs derived from HepG2 and Huh7 showed significant dissimilarity (Figure 10 a, m^1^A logos). The Tomtom similarity heatmap for m^1^A (Figure 10c, left panel) displays overall low − log_10_(*q*-value) scores, primarily ranging below 1.5 (e.g., maximum score of 1.8), confirming that the local sequence context recognized by EvoRMD for m^1^A prediction is highly cell-type-specific. To further quantify this divergence, we performed Principal Component Analysis (PCA) on the 3-mer frequency composition of the extracted motifs. Consistent with the motif logos and similarity matrix, the PCA results for m^1^A show that motifs from HepG2 and Huh7 form two distinct, well-separated clusters, indicating fundamentally different underlying sequence features captured by the model in the two cell lines (Figure 10 b, m^1^A PCA), indicating fundamentally different underlying sequence features captured by the model in the two cell lines. This divergence is fundamentally rooted in the structure-dependent nature of m^1^A writers (e.g., TRMT6*/*TRMT61A), which primarily recognize RNA secondary structure (such as stem-loops) and local folding state, contrasting sharply with the sequence-driven recognition of m^6^A ^70,71^. The resulting m^1^A epitranscriptomic landscape is highly plastic and susceptible to cell-specific contextual factors, including heterogeneous m^1^A writer expression levels, distinct UTR folding patterns influenced by the two cell lines’ unique metabolic/phenotypic backgrounds, and the resulting enrichment of distinct target RNA clusters. This profound dependence on structural and contextual cues causes m^1^A to be deposited in completely different sequence backgrounds, leading to motifs that are fundamentally dissimilar and providing strong evidence that EvoRMD successfully resolves this biological plasticity.

**Fig. 10.**
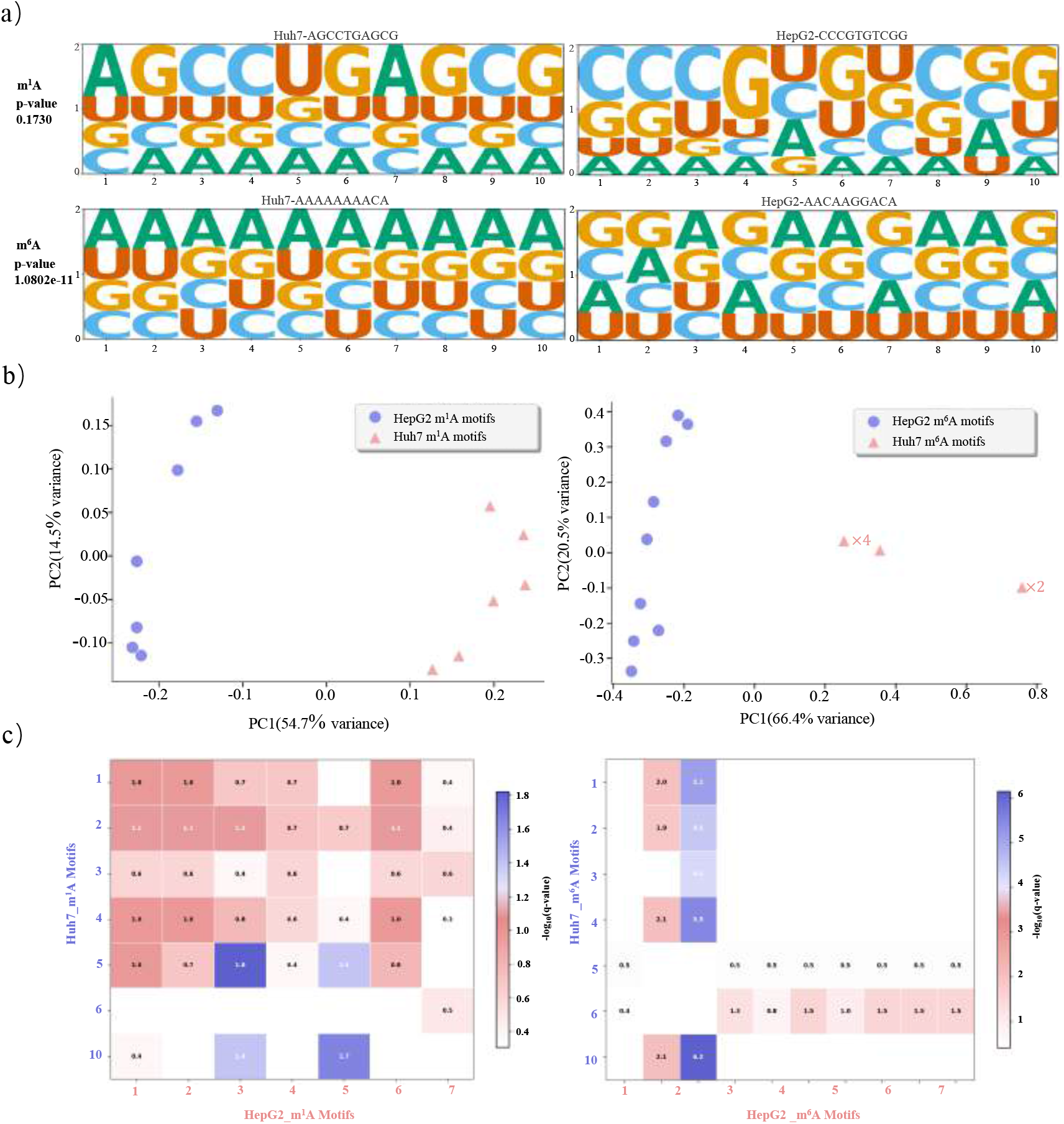
Cell-type-specific motif divergence and conservation analysis in Hepatocellular Carcinoma (HepG2 vs Huh7) cell lines. **a)** Sequence logos derived from EvoRMD attention-enriched windows. The m^1^A motifs (top row) show high divergence between HepG2 and Huh7, while m^6^A motifs (bottom row) exhibit high conservation, converging toward the established RRACH canonical consensus. **b)** Principal Component Analysis (PCA) of 3-mer frequency composition for extracted motifs. m^1^A motifs (left) form two distinct, well-separated clusters, indicating fundamentally different underlying sequence features captured by the model. m^6^A motifs (right) show structured separation, reflecting subtle differences in flanking sequence composition despite core conservation.**c)** Tomtom cross-cell line similarity heatmap ( − log_10_(*q-value*)). The m^1^A heatmap (left) displays overall low scores (darker blocks, max ≈ 1.8), confirming motif dissimilarity and cell-type specificity. The m^6^A heatmap (right) shows significantly high scores (brighter blocks, max 6.2), demonstrating strong motif conservation.

In contrast, the m^6^A motifs exhibited remarkable conservation between HepG2 and Huh7. The motif logos for m^6^A (Figure 10 a, m^6^A logos) are highly similar, converging onto the established RRACH canonical consensus ^72^. The Tomtom similarity heatmap for m^6^A (Figure 10 c, right panel) shows a drastically different pattern, featuring high − log_10_(*q*-value) scores, with several strong matches exceeding 5.0 (maximum score of 6.2). These high scores, particularly for key Huh7 motifs matching specific HepG2 motifs (e.g., Huh7 10 vs. HepG2 3, score 6.2), demonstrate strong cross–cell-line matching and a highly conserved sequence signal. The 3-mer PCA analysis for m^6^A further confirms this conservation, yet reveals a structured separation in the reduced-dimensional space (Figure 10 b, m^6^A PCA), with HepG2 motifs generally clustering separately from Huh7 motifs along PC1. We interpret this structured separation as reflecting subtle, consistent differences in the flanking sequence composition that EvoRMD utilizes as cellline-specific contextual features, even as the core m^6^A recognition motif remains functionally conserved, as evidenced by the high Tomtom scores.

The stark contrast between m^1^A divergence and m^6^A conservation within the same pair of cell lines highlights a crucial biological insight captured by EvoRMD: m^6^A is likely governed by a more rigid, sequence-dependent core recognition complex, while the regulatory context for m^1^A is more plastic and susceptible to cell-state-specific regulatory factors, leading to the evolution of distinct epitranscriptomic signatures. This ability to resolve both conserved and divergent motifs for different modification types is strong evidence that EvoRMD is learning true biological signal, rather than merely recognizing global sequence biases.

#### 2.5.2 Malignant Transformation (HNPCs vs GSCs)

To further investigate the impact of malignant transformation on epitranscriptomic regulation, we extended the analysis to compare the m^6^A motifs shared between human neural progenitor cells (HNPCs) and highly malignant glioma stem cells (GSCs) (Figure 11). The Tomtom similarity heatmap (Figure 11 c) and motif logos (Figure 11 a) initially confirm that the core m^6^A recognition motif remains highly conserved across the HNPC-GSC transformation, supported by high cross-cell line Tomtom matching scores and the extremely low *p*-value (e.g., *p* ≈ 4.7 *×* 10^−10^) for the strongest motif match, which confirms the integrity of the canonical RRACH core signal. However, this conservation of the enzymatic recognition core contrasts sharply with a striking cell-type–specific divergence in the overall motif environment. The PCA plot (Figure 11 b, left) demonstrates a significant distinction based on motif 3-mer composition, with PC1 explaining a substantial 76.7% of the variance, confirming that EvoRMD captures highly divergent local sequence features. This divergence is entirely driven by the flanking sequence composition, as visualized in the composition scatter plot (Figure 11 b, right).

**Fig. 11.**
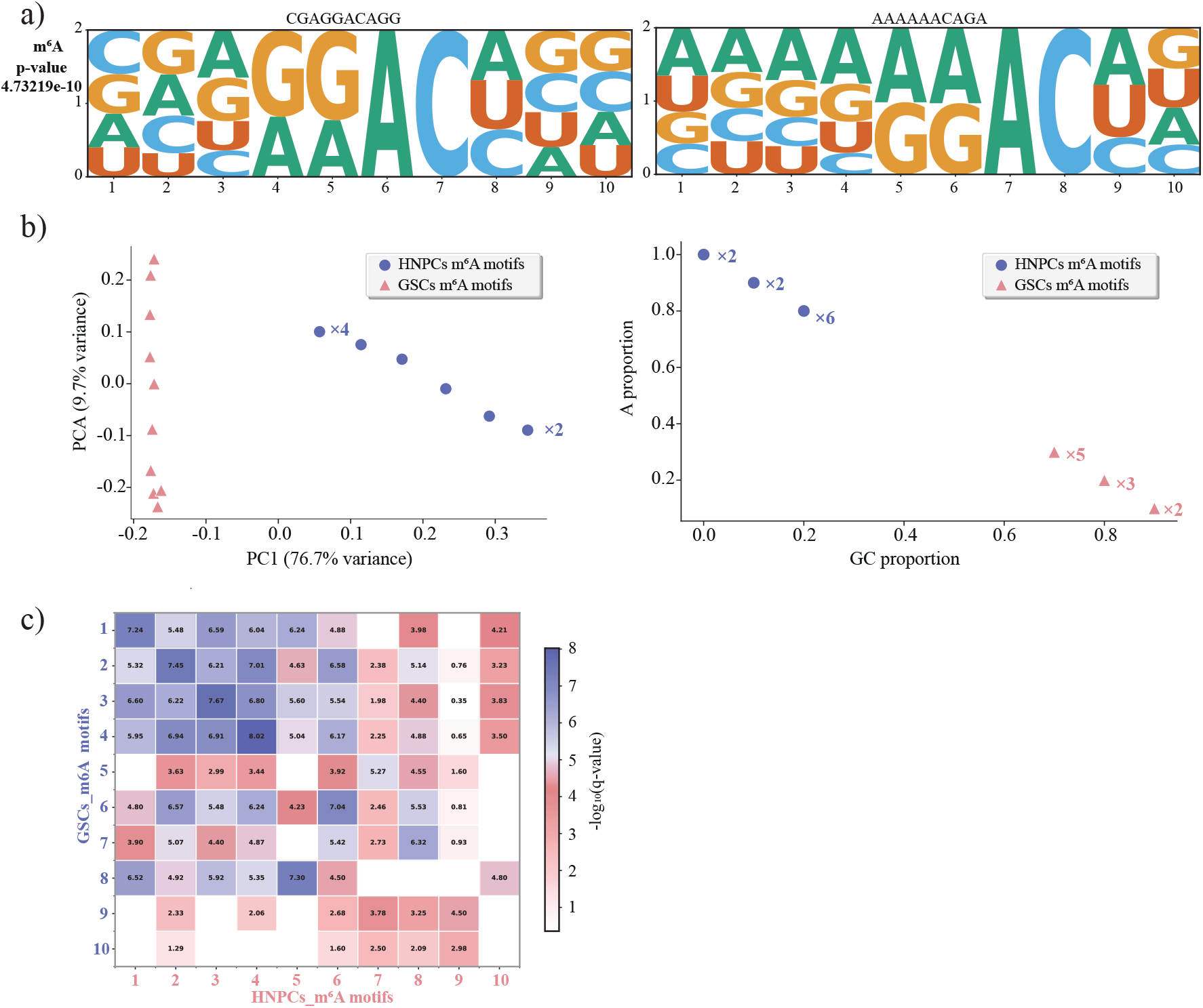
Cell-type-specific motif signature divergence in Neural Lineage (HNPCs vs GSCs). **a)** Sequence logos derived from EvoRMD attention-enriched windows for m^6^A motifs in GSCs and HNPCs. Both motifs contain the established RRACH canonical core (e.g., ACAG/CAG), but the flanking sequences exhibit high divergence. **b)** Principal Component Analysis (PCA) and sequence composition divergence. The PCA plot (left) shows that m^6^A motifs from GSCs and HNPCs form two distinct, well-separated clusters (PC1 explains 76.7% of the variance), indicating fundamentally different underlying sequence features in the local context. The scatter plot (right) visualizes the divergence: GSC motifs cluster tightly in the GC-rich region (high GC proportion, low A proportion), while HNPC motifs show a strong enrichment in the A-rich region (high A proportion, low GC proportion). **c)** Tomtom cross-cell line similarity heatmap ( − log_10_(*q-value*)). The heatmap displays overall low to moderate scores (predominantly in the 2.0-5.0 range), confirming that while the canonical core is shared, the broader 10bp motif context is dissimilar between the cell lines.

A closer analysis of the sequence context reveals a profound shift in regulatory preference associated with malignant transformation. Motifs derived from GSCs display a strong preference for a GC-rich, G-rich flanking environment (e.g., CGGACAGGGG), as confirmed by their tight clustering in the high GC proportion/low A proportion region of the composition scatter plot (Figure 11 b, right). This GSC-specific bias suggests that the methylation writers may preferentially target m^6^A sites within contexts prone to secondary structure or associated with actively regulated genomic regions. Conversely, HNPC motifs are strongly characterized by A-rich, polyA-like flanking regions (e.g., AAAAAACAGA), leading to their distinct clustering in the high A proportion/low GC proportion region (Figure 11 b, right). This cell-type-specific m^6^A motif signature, defined by divergent flanking contexts around a conserved core, suggests that m^6^A regulation in normal progenitor cells is associated with a distinct set of transcripts or structural contexts, possibly linked to differential UTR folding or stability mechanisms. This ability of EvoRMD to capture a shift in the local sequence context, even for a core-conserved modification, underscores its power to resolve biologically meaningful contextual regulation driven by malignant transformation.

#### 2.5.3 Functional consequences of cell-type specific RNA modification patterns

To address whether the distinct sequence motifs observed above translate into broader biological differences, we analyzed the genomic distribution and functional enrichment of modification sites unique to each cell type. We first examined the overlap of modified genes between cell lines. As shown in Figure 12 b), the overlap of *m*^6^*A* targets between HepG2 and Huh7 and between HNPCs and GSCs is remarkably low, confirming that distinct cell states utilize the epitranscriptome to regulate largely disjoint sets of transcripts. Furthermore, the genomic distribution of these sites (Figure 12 a)) reveals subtle but specific shifts; for instance, *m*^1^*A* sites in HepG2 show a distinct preference for CDS regions compared to Huh7, reflecting the structural plasticity of *m*^1^*A* regulation identified in our motif analysis.

**Fig. 12.**
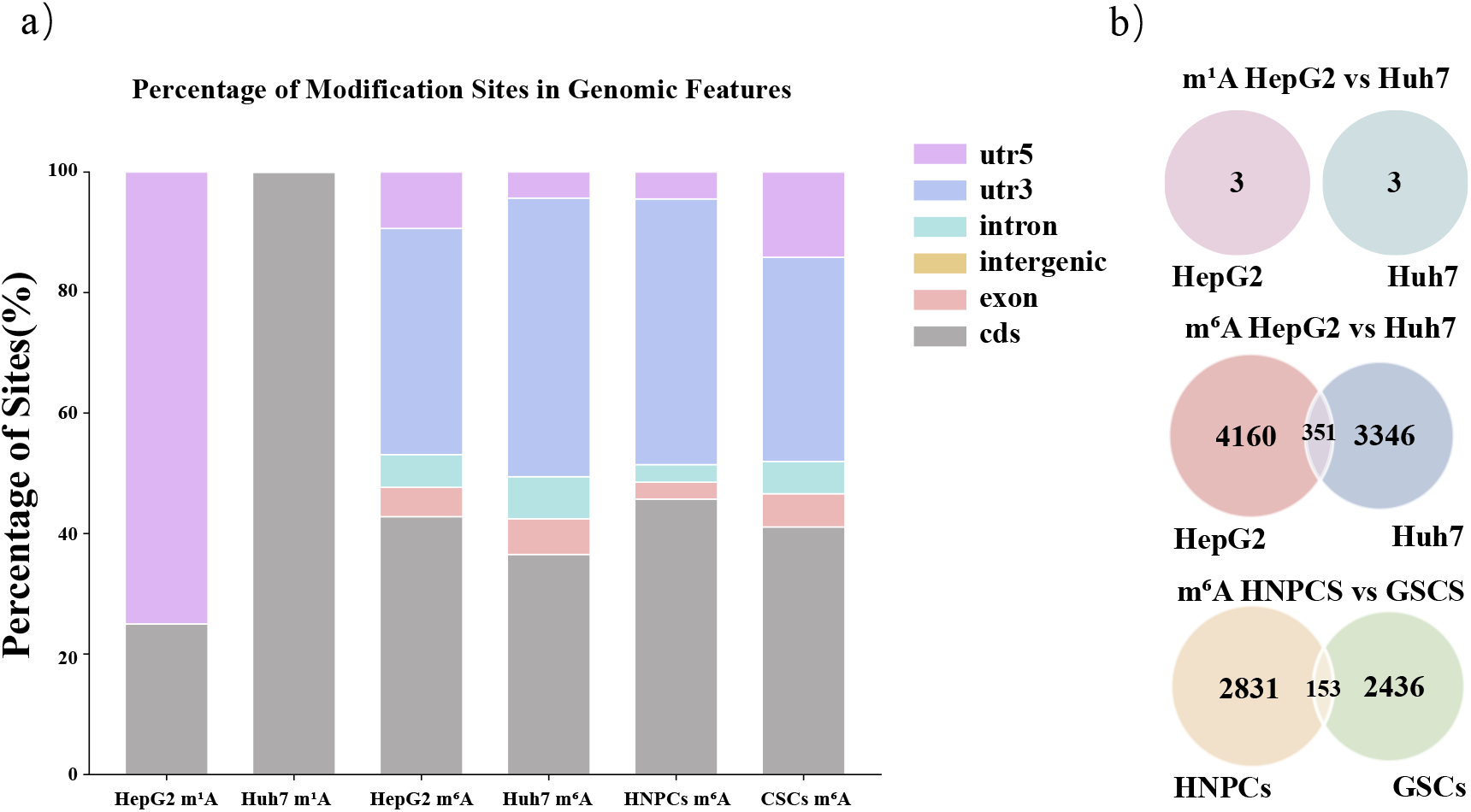
Broader functional characterization of cell-type specific RNA modification profiles. **a)** Genomic distribution of modification sites. The distribution across transcript regions (e.g., 5’UTR, CDS) reveals cell-type-specific patterns (e.g., *m*^1^*A*) that mirror the observed motif plasticity. **b)** Target gene overlap. Venn diagrams show limited overlap of modified genes between cell lines, indicating highly distinct epitranscriptomic landscapes.

Crucially, functional enrichment analysis of target genes associated with EvoRMD-identified cell-type-specific motifs using the Reactome ^73^ database reveals that EvoRMD captures biologically coherent regulatory programs consistent with the specific phenotype of each cell line (Figure 13). In Hepatocellular Carcinoma (HepG2 vs. Huh7), HepG2-specific *m*^6^*A* targets are significantly enriched in pathways related to “TP53 Regulates Transcription of Genes Involved in Cytochrome C Release” and “Sphingolipid Metabolism”. This is biologically consistent with the genetic background of HepG2 cells, which possess wild-type p53 capable of initiating apoptosis via the cytochrome C pathway ^74,75^, and reflects their active role in hepatic lipid homeostasis ^66^. In contrast, Huh7-specific targets are notably enriched in “Glucuronidation” and “Metabolism of proteins” related pathways. The enrichment of glucuronidation—a key phase II detoxification process—aligns with the extensive use of Huh7 models in drug metabolism (ADME) and toxicology studies due to their distinct metabolic enzyme profiles compared to HepG2^60,76^. In Neural Lineage Transformation (HNPCs vs. GSCs), the transition from normal progenitors to cancer stem cells is marked by a dramatic functional shift. HNPC-specific *m*^6^*A* targets are heavily enriched in “Translation”, “Peptide Chain Elongation”, and “Selenocysteine Synthesis”, supporting the high protein synthesis demands required for the rapid proliferation and maintenance of normal progenitor cells ^77^. Conversely, GSC-specific targets are enriched in “FRS-mediated FGFR4 Signaling” and “Defects in Vitamin and Cofactor Metabolism”. The appearance of FGFR4 signaling is particularly relevant, as fibroblast growth factor receptors are known drivers of tumorigenesis and stemness in glioblastoma ^78**?**^ . Additionally, the identification of “Phase 4 - Resting Membrane Potential” pathways in GSCs correctly identifies the retained neural electrophysiological signature that drives glioma progression and integration into neural circuits ^79^.

**Fig. 13.**
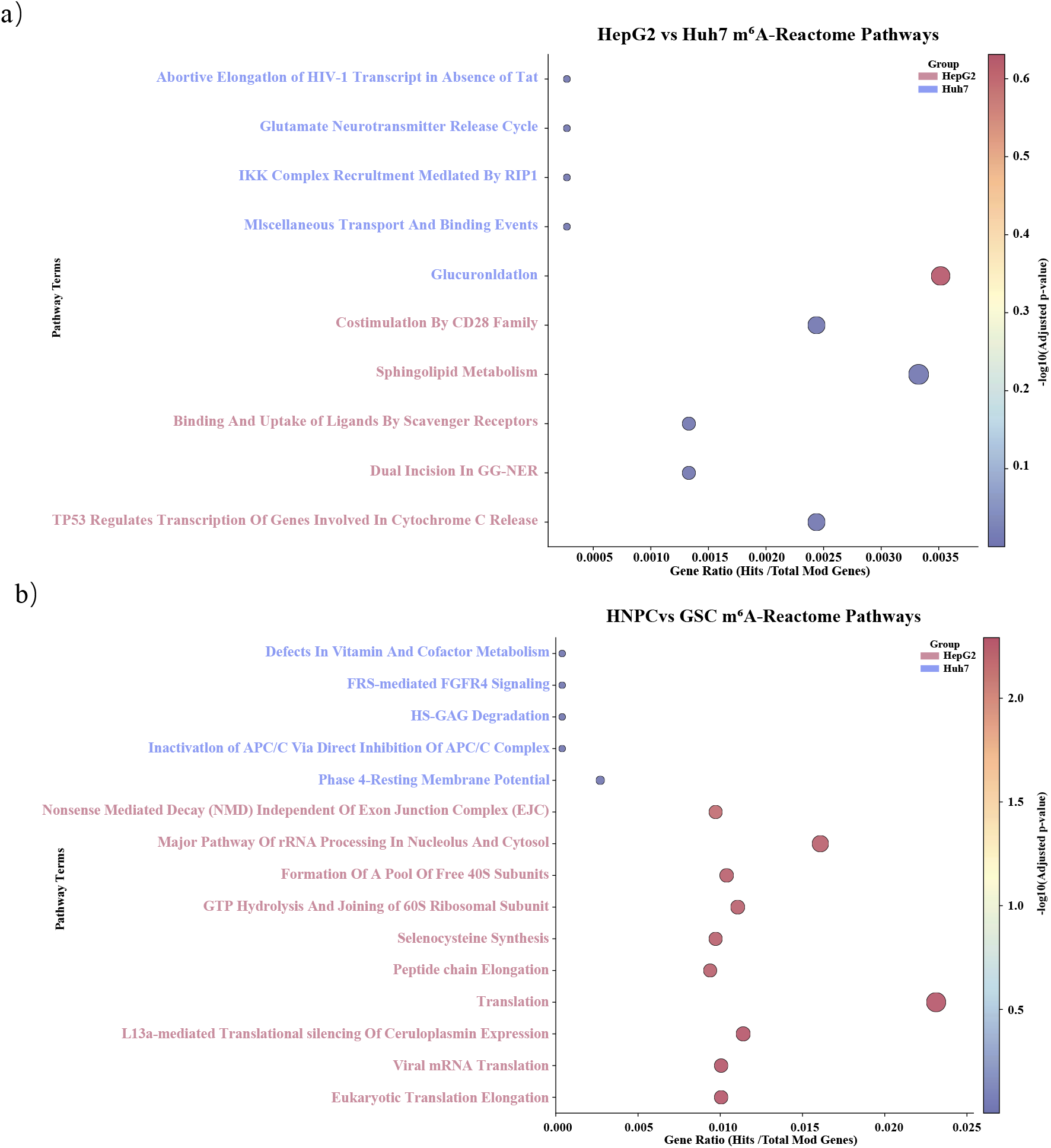
Functional enrichment analysis of cell-type specific *m*^6^*A* regulatory programs. The scatter plots visualize enriched Reactome pathways, where the x-axis represents the Gene Ratio, bubble size indicates gene count, and color intensity reflects statistical significance ( − log_10_(Adjusted P-value)). **a)** HepG2 vs. Huh7: HepG2 targets (pink) are significantly enriched in sphingolipid metabolism and p53-mediated cytochrome C release, while Huh7 targets (blue) are characterized by glucuronidation and drug metabolism (ADME) pathways. **b)** HNPC targets (pink) are dominated by translation and peptide elongation pathways supporting rapid proliferation, whereas GSC targets (blue) highlight FGFR4 signaling, vitamin/cofactor metabolic defects, and resting membrane potential, reflecting the unique metabolic and stemness features of glioblastoma.

In summary, the distinct motifs identified by EvoRMD are not stochastic variations but are linked to specific, functionally coherent gene regulatory networks that drive cell identity and disease states.

### 2.6 Ablation study

#### 2.6.1 Downsampling for class balance

To reduce the problem of class imbalance while maintaining diversity among species, we downsampled the large majority modifications (m^6^A and m^5^C) and retained all samples of the other modifications. For each target modification with original total *n*_total_, we set its downsampled total as

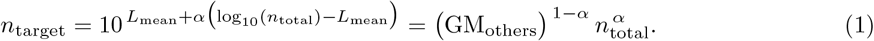

Here, *L*_mean_ is the base-10 logarithm of the geometric mean of the totals of the other (non-target) modifications, i.e.. The parameter *α* ∈ [0, 1] controls the compression strength; smaller *α* implies stronger compression. When *α* = 0, the target size collapses to the geometric-mean scale of the other (non-target) modifications.

We aggregate available samples by species and use log(available+1) as allocation weights to distribute the target total *n*_target_ across species. For each species, we enforce a minimum keep (default min keep = 1) and cap the quota by its actual availability. If a *remainder* persists after rounding, we apply a greedy refill ordered by the remaining capacity. The samples in the non-target modification category are not downsampled.

We compared *α* ∈ *{*0, 0.2, 0.4, 0.6, 0.8, 1.0*}* using three random seeds (42, 215, 3407). As shown in Table 6, the model consistently achieved high per-class and overall MCC scores across all downsampling strengths. Nevertheless, combining both performance and training efficiency (i.e., faster convergence), *α* = 0.6 provided the most favorable balance (Table 6 and Fig. 14). Therefore, all reported experiments are based on *α* = 0.6.

**Table 6.**
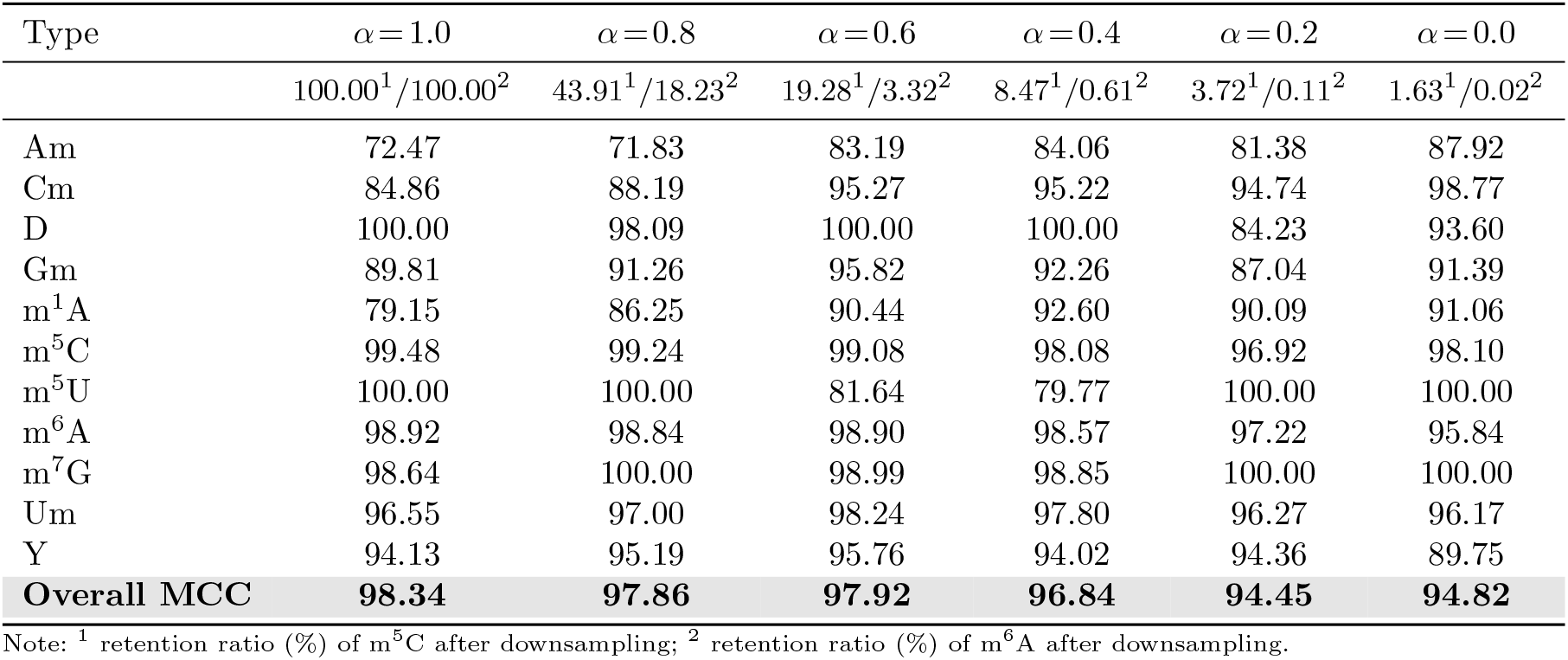
Per-class MCC (%) across different *α* values (averaged over three seeds).

| Type | $\alpha=1.0$ | $\alpha=0.8$ | $\alpha=0.6$ | $\alpha=0.4$ | $\alpha=0.2$ | $\alpha=0.0$ |
| --- | --- | --- | --- | --- | --- | --- |
|  | 100.00<sup>1</sup>/100.00<sup>2</sup> | 43.91<sup>1</sup>/18.23<sup>2</sup> | 19.28<sup>1</sup>/3.32<sup>2</sup> | 8.47<sup>1</sup>/0.61<sup>2</sup> | 3.72<sup>1</sup>/0.11<sup>2</sup> | 1.63<sup>1</sup>/0.02<sup>2</sup> |
| Am | 72.47 | 71.83 | 83.19 | 84.06 | 81.38 | 87.92 |
| Cm | 84.86 | 88.19 | 95.27 | 95.22 | 94.74 | 98.77 |
| D | 100.00 | 98.09 | 100.00 | 100.00 | 84.23 | 93.60 |
| Gm | 89.81 | 91.26 | 95.82 | 92.26 | 87.04 | 91.39 |
| m<sup>1</sup>A | 79.15 | 86.25 | 90.44 | 92.60 | 90.09 | 91.06 |
| m<sup>5</sup>C | 99.48 | 99.24 | 99.08 | 98.08 | 96.92 | 98.10 |
| m<sup>5</sup>U | 100.00 | 100.00 | 81.64 | 79.77 | 100.00 | 100.00 |
| m<sup>6</sup>A | 98.92 | 98.84 | 98.90 | 98.57 | 97.22 | 95.84 |
| m<sup>7</sup>G | 98.64 | 100.00 | 98.99 | 98.85 | 100.00 | 100.00 |
| Um | 96.55 | 97.00 | 98.24 | 97.80 | 96.27 | 96.17 |
| Y | 94.13 | 95.19 | 95.76 | 94.02 | 94.36 | 89.75 |
| **Overall MCC** | **98.34** | **97.86** | **97.92** | **96.84** | **94.45** | **94.82** |

Note: <sup>1</sup> retention ratio (%) of m<sup>5</sup>C after downsampling; <sup>2</sup> retention ratio (%) of m<sup>6</sup>A after downsampling.

**Fig. 14.**
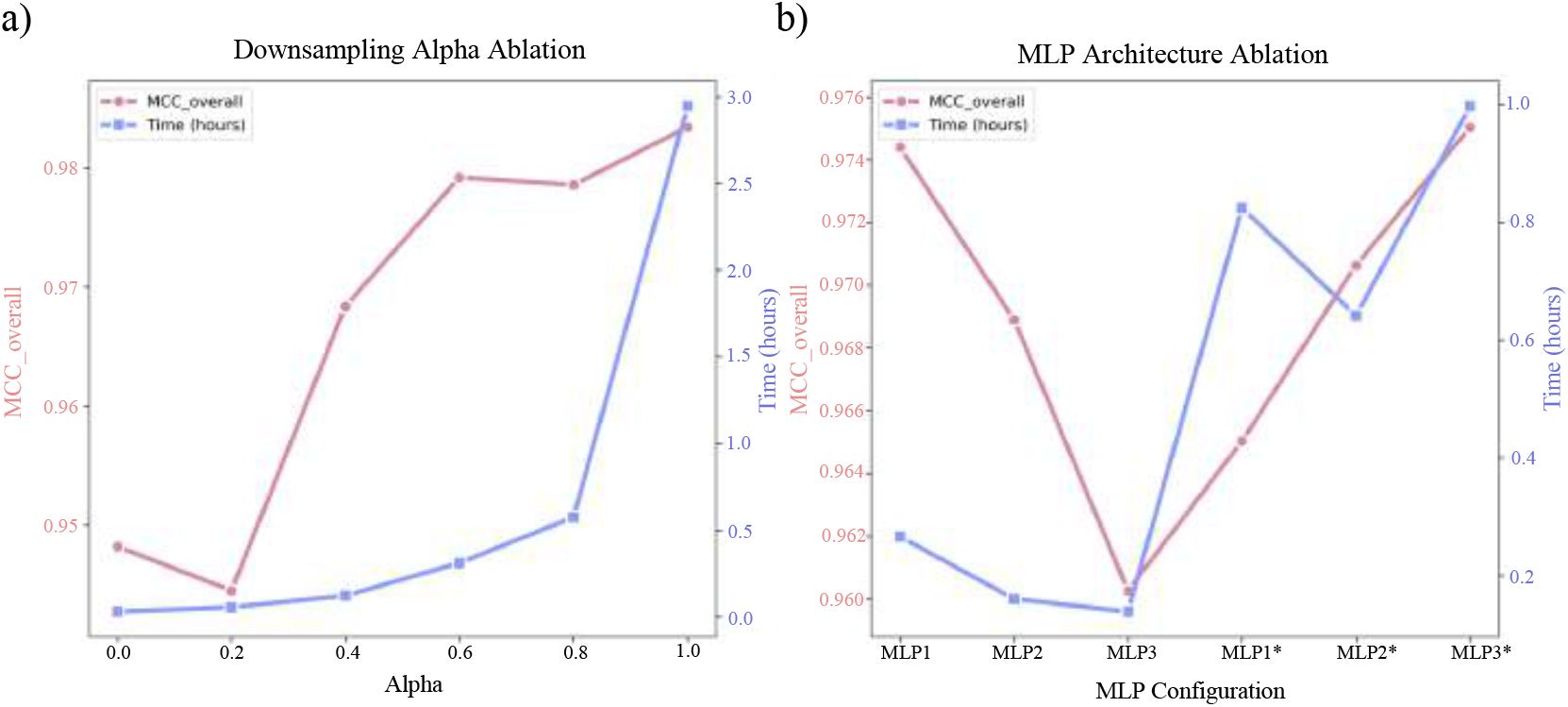
Performance and training-time ablation results. **a)** Effect of downsampling strength (*α*) on model performance and convergence efficiency. **b)** Impact of MLP depth and RNA-FM fine-tuning on overall MCC and training cost. Together, these analyses identify *α* = 0.6 and a three-layer fine-tuned MLP (MLP3*) as the most effective configuration.

#### 2.6.2 Classifier depth and RNA-FM fine-tuning

To evaluate how architectural choices influence EvoRMD’s performance, we conducted an ablation study examining two factors: (i) the depth of the classification head and (ii) whether the RNA-FM encoder is kept frozen or jointly fine-tuned. All experiments were run with three random seeds (42, 215, 3407), and performance metrics represent the mean across runs. Table 7 summarizes the mean MCC and time-to-best epoch for each configuration.

**Table 7.** Mean MCC (%) and mean time to best epoch (minutes) over three seeds (42, 215, 3407) for different MLP depths with and without RNA-FM fine-tuning (Downsampling *α* = 0.6).

| Classifier depth | Not fine-tuned |  | Fine-tuned |  |
| --- | --- | --- | --- | --- |
|  | Mean MCC (%) | Time to best (min) | Mean MCC (%) | Time to best (min) |
| Shallow (0 hidden layer) | **97.44** | 15.96 | 96.50 | 49.44 |
| Moderately deep (1 hidden layer) | 96.89 | 9.61 | 97.06 | 38.47 |
| Deep (2 hidden layers) | 96.03 | 8.31 | **97.50** | 59.82 |

Note: **Bold** numbers indicate the best performance within each column (highest MCC).

**Table 8:** Details of the 11 RNA modification datasets.

| Type | Am | Cm | Gm | Um | D | Y | m<sup>1</sup>A | m<sup>5</sup>C | m<sup>5</sup>U | m<sup>6</sup>A | m<sup>7</sup>G |
| --- | --- | --- | --- | --- | --- | --- | --- | --- | --- | --- | --- |
| Nucleotide | A | C | G | U | U | U | A | C | U | A | G |
| Count | 426 | 717 | 301 | 1,579 | 96 | 277 | 842 | 21,097 | 27 | 1,713,087 | 778 |
| Downsampled Count* | – | – | – | – | – | – | – | 4,068 | – | 56,901 | – |

Note: \* Values represent the downsampled dataset sizes obtained under the downsampling parameter $\alpha = 0.6$ . This setting achieves the best trade-off between overall model performance and convergence speed. All model training uses the $\alpha = 0.6$ downsampled data. Details can be found in the ablation study section.

We compared three classifier designs spanning a practical range of capacities used in sequence-based prediction tasks: a *shallow* classifier consisting of a single linear projection (no hidden layers), a *moderately deep* classifier with one hidden layer and nonlinear activation, and a *deep* classifier comprising two hidden layers. When the RNA-FM encoder is frozen, increasing classifier depth does not produce consistent gains; in fact, the shallow classifier achieves the highest mean MCC (97.44%), whereas deeper variants show mild performance declines, indicating that frozen RNA-FM representations are already sufficiently informative for linear separation.

Fine-tuning RNA-FM substantially changes this pattern. Once the encoder is trainable, deeper classifiers more effectively leverage the richer, co-adapted sequence–context representations and achieve higher performance, with the deep classifier reaching the highest overall MCC (97.50%). Although fine-tuning introduces additional computational cost that scales with classifier depth (Fig. 14), the total overhead remains modest (≤1 hour). Overall, these results demonstrate a clear interaction between encoder trainability and classifier capacity. While a shallow head suffices when RNA-FM is frozen, the combination of a deep classifier and a fine-tunable RNA-FM encoder provides the best balance of predictive accuracy and computational efficiency, and is therefore adopted as the default EvoRMD configuration for all subsequent analyses.

## 3 Discussion

The EvoRMD model introduces a unified and biologically grounded framework that significantly advances RNA modification prediction. By integrating evolutionary-aware embeddings from RNA-FM ^34^ with a hierarchical anatomical encoder that jointly models species, organ/tissue or cell line, and subcellular localization, EvoRMD captures both sequence-level and contextual determinants of RNA modifications. Together with a specialized attention mechanism and a unified multi-class architecture, this design yields substantial improvements over existing multi-modification predictors while remaining computationally efficient. The attention module dynamically weighs nucleotide positions within their biological context, enabling the model to highlight sequence features that are likely involved in recognition by modification-related enzymes or RNA-binding proteins. Beyond accurate classification, EvoRMD recovers conserved sequence motifs that align well with known biological motifs recorded in the RMBase database ^18^, providing strong evidence for the biological validity of its internal representations. Moreover, by analyzing EvoRMD-derived motifs across different cell types and states (e.g., HepG2 vs Huh7, HNPCs vs GSCs), the model reveals both conserved and cell-specific motif variants, as well as notable correlations among different RNA modification types that suggest potential crosstalk or coordinated regulation mediated by shared enzymatic pathways or modification hierarchies. These insights open avenues for a deeper understanding of epitranscriptomic regulatory networks in their native biological context.

Despite these advances, several limitations warrant further exploration. Although EvoRMD now achieves strong performance even for nominally rare modifications such as Cm and m^5^U, the absolute number of experimentally labeled sites for these categories remains very small, which calls for cautious interpretation and highlights the need for larger and more diverse datasets. In this context, combining EvoRMD with explicit few-shot or meta-learning and semi-supervised strategies for ultra-rare or newly profiled modifications is an important direction for future work as additional data become available. In addition, while EvoRMD provides rich computational evidence through attention patterns, cell-specific motif discovery, and co-modification analyses, detailed biochemical and structural experiments remain essential to validate individual predictions and to confirm that the inferred sequence and context signatures reflect genuine mechanisms rather than residual dataset or model biases. Finally, although our analyses across multiple cell types and malignant vs non-malignant states demonstrate the potential of EvoRMD for biological and disease-related applications, systematic integration with patient cohorts and clinical phenotypes lies beyond the scope of the present study. Future work should therefore focus on (i) coupling EvoRMD with experimental validation pipelines, (ii) extending it with advanced learning strategies tailored to ultra-rare and newly discovered modifications, and (iii) applying it to large-scale disease association, biomarker discovery, and therapeutic target prioritization to further realize its translational potential.

## 4 Conclusions

In conclusion, EvoRMD offers a robust and interpretable computational framework for predicting multiple RNA modification types from RNA sequences. By leveraging an evolutionary-aware RNA language model together with a hierarchical anatomical encoder that integrates species, organ/tissue, cell line, and subcellular localization, EvoRMD captures both sequence and biological-context signals and achieves state-of-the-art performance across diverse RNA modifications. The unified multi-class architecture further simplifies multi-modification prediction, enhancing computational efficiency and scalability. Importantly, the attention mechanism in EvoRMD not only highlights nucleotide positions that are crucial for prediction, but also enables the discovery of biologically meaningful conserved motifs and their cell-specific variants. By comparing motifs across different cell types and states (such as HepG2 vs Huh7 and HNPCs vs GSCs), EvoRMD reveals where modification signals are evolutionarily conserved and where they become cell- or state-dependent. Thus, EvoRMD not only improves predictive accuracy, but also provides biologically interpretable insights into the context-dependent regulation of RNA modifications, laying a solid foundation for future studies on RNA epitranscriptomic mechanisms and their roles in broader biological and clinical settings.

## 5 Methods

### 5.1 Problem formulation

Current epitranscriptomic profiling assays are modification-specific—such as antibody enrichment, chemical conversion, or mutation-based detection—and therefore identify only a single modification type at each genomic coordinate. Thus, while one modification label is observed for each site, all other types remain *unobserved* and cannot be treated as true negatives. This produces a single-positive, multiple-unlabeled (SP–MU) supervision regime, a standard form of partial-label learning.

For each candidate RNA site, the model observes a 41-nt sequence window **x** and structured biological content **c** = *{c*_*o*_, *c*_*ct*_, *c*_*sl*_, *c*_*sp*_*}*, describing the organ, cell type, subcellular compartment, and species. Each site is annotated with exactly one observed modification type, *y* ∈ *{*1,. .., *K}*, where *K* denotes the total number of candidate RNA modification classes. Given that only a single observed label is available per site, the supervised task corresponds to a single-label, multi-class classification problem.

EvoRMD encodes the multimodal input (**x, c**) into a unified representation **h** ∈ ℝ^*d*^, which is passed to a classification head producing logits **z** = *f*_*θ*_(**h**) ∈ ℝ^*K*^ . Because the SP–MU setting does not justify treating unobserved types as negatives, these logits are interpreted through a softmax distribution:

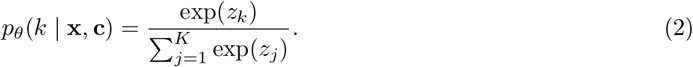

This formulation yields a context-dependent ranking over all modification types, reflecting the most plausible outcome without imposing negative assumptions on unobserved labels.

### 5.2 Training objective

As each RNA site provides only a single observed label under the SP–MU regime, EvoRMD is trained using a standard single-label, multi-class objective implemented via softmax cross-entropy. The model is encouraged to assign high probability to the observed modification *y*_*i*_ (with *y*_*i*_ ∈ *{*1,…, *K}* indexing the *K* candidate modification types) while making no assumptions about the unobserved types.

Given the softmax-normalized probabilities *p*_*θ*_(*k* | **x, c**), the training objective is:

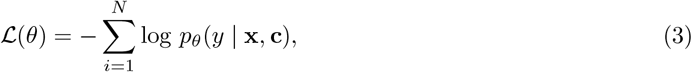

where *N* is the number of training examples. Because unobserved modification types are not treated as negatives, the loss does not penalize the model for assigning nonzero probability to them. Instead, the softmax distribution provides a context-dependent ranking over the *K* candidate classes.

### 5.3 Post-hoc Multi-Label Inference

Although EvoRMD is trained with a single-label softmax objective, it outputs a logit *z*_*k*_ for every modification type *k*. To enable fair comparison with binary and multi-label predictors, we convert these logits into independent likelihoods using a sigmoid transformation, *p*_*k*_ = *σ*(*z*_*k*_), and apply per-class decision thresholds.

Because unobserved modification types cannot be treated as negatives in the SP–MU regime, threshold selection cannot rely on raw class imbalance. For each modification type *k*, we construct a balanced validation subset with a 1:10 positive-to-sampled-negative ratio and select the threshold 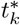 maximizing the Matthews correlation coefficient (MCC):

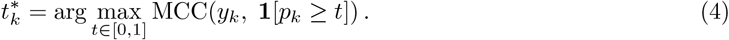

At test time, EvoRMD produces a multi-label output via 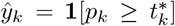. This procedure yields independent predictions for each modification type without retraining the model and enables direct, equitable comparison with binary or multi-label baselines while respecting the SP–MU constraints.

### 5.4 Architecture

#### 5.4.1 Encoder module

For each sample, the encoder takes as input a RNA sequence **x** and structured biological content **c** = *{c*_*o*_, *c*_*ct*_, *c*_*sl*_, *c*_*sp*_*}*, representing the organ, cell type, subcellular location, and species.

##### Embedding of biological context

Each metadata field is first mapped to a hybrid multi-hot vector, which assigns in-vocabulary labels to dedicated indices, routes out-of-vocabulary entries to *H* hash buckets (default *H* = 256 per level), and uses a field-specific “unknown” token for missing values. This strategy preserves expressiveness, supports unseen labels during inference, and prevents uncontrolled growth in parameter dimensionality. For each metadata component *m* ∈ *{o, ct, sl, sp}*, we obtain a dense embedding via a category-specific Linear–LayerNorm transformation:

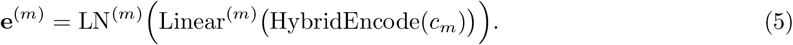

To capture hierarchical biological structure, we average the organ, cell-type, and subcellular-location embeddings,

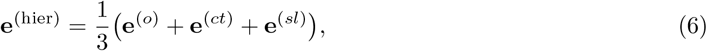

and concatenate this with the species embedding to obtain the final context representation:

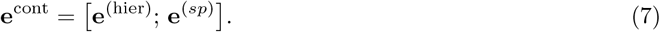

##### Embedding of RNA sequence

For each sample, we use a 41-nt sequence centered on the candidate modification as the model input. RNA-FM ^34^, a pretrained RNA foundation model, produces contextualized token embeddings

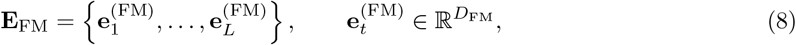

so that 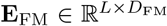, where *L* denotes the number of tokens (41 nucleotides plus two special tokens [`CLS`] and [`SEP`], giving *L* = 43 in our implementation) and *D*_FM_ is the dimensionality of the RNA-FM token embeddings. In our experiments, we use the hidden states from the final (12th) transformer layer of RNA-FM as the sequence features.

##### Concatenated Embedding

To condition each nucleotide on its biological context, the metadata embedding is broadcast along the sequence length,

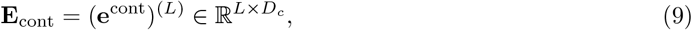

and concatenated with the RNA-FM embeddings at every position:

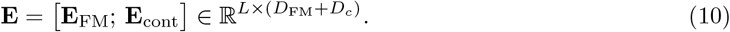

The matrix **E** serves as the multimodal encoder output for each sequence, jointly capturing sequence-derived and context-derived information.

#### 5.4.2 Attention module

To obtain a site-level representation from the position-wise encoder output, EvoRMD applies a trainable attention mechanism. Each token embedding **e**_*i*_ (the *i*-th row of **E**) is assigned an attention weight

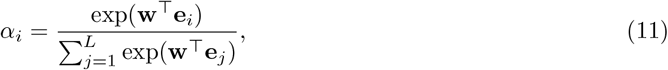

where **w** is a learned parameter vector. The site-level representation is then computed as an attention-weighted sum:

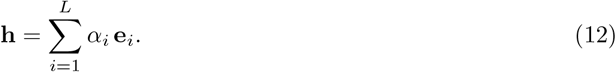

This aggregation allows the model to focus on positions and sequence–context combinations that are most informative for predicting the modification type.

#### 5.4.3 Classifier module

The site-level representation **h** is passed through a three-layer multilayer perceptron (MLP) to produce logits over the *K* modification classes. The MLP consists of two hidden layers with ReLU activations followed by a linear output layer:

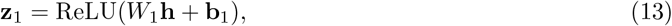

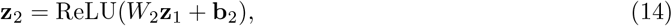

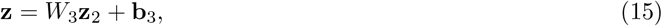

where **z** ∈ ℝ^*K*^ are the final logits used in the softmax-based single-label, multi-class objective described in Sections 5.1 and 5.2. This architecture enables EvoRMD to learn discriminative site-level features that integrate both sequence signals and biological context to predict RNA modification types.

### 5.5 Experiments setting

#### 5.5.1 Data preparation

We assembled experimentally supported sites for 11 RNA modifications (Am, Cm, Gm, Um, D, Y, m^1^A, m^5^C, m^5^U, m^6^A, m^7^G) from RMBase v3.0^18^ across 10 species (*Homo sapiens* hg38, *Mus musculus* mm10, *Rattus norvegicus* rn6, *Oryctolagus cuniculus* OryCun2, *Ovis aries* Oarv3, *Mesocricetus auratus* MesAur1, *Bos taurus* bosTau9, *Macaca mulatta* rheMac8, *Pan troglodytes* panTro5, *Sus scrofa* susScr11). For each site, we extracted a 41-nt sequence window (*±*20 nt) centered on the modification coordinate. To enable downstream encoding as a structured RNA-context modality embedding, we normalized the RMBase field cellList (the list of cells or tissues in which this RNA modification is identified) into three orthogonal columns: `cell` (cell line or primary cell; e.g., HEK293T→HEK293, GM12878/LCL→LCLs, ESC/mESC/hESC→ESCs),`organ` (e.g., liver, lung, brain, lymphatic system), and `subcellular location` (keyword mapping such as cytosolic→cytoplasm, mitochondrial→mitochondria, nucleoplasmic→nucleus). The mouse strain token “C57BL6” denotes genetic background rather than a cell or tissue: rows containing only “C57BL6” were removed; for multi-token entries, only this token was dropped and the remaining valid tokens were retained. To control redundancy, 41-nt windows were de-duplicated using CD-HIT ^80,81^ at 80% sequence identity (`-c 0.8 -n 5 -g 1 -d 0`). Unlike global clustering, we performed CD-HIT separately for each combination of species and modification type, ensuring that sequence diversity was preserved within biological categories. Within each (species, modification) cluster, one representative sequence was retained with the fewest “unknown” values across (`organ`, `subcellular_location`, `cell`), and in case of ties, retained the record with the smallest original index.

Compared with conventional binary classification frameworks that rely on artificially constructed positive and negative samples, EvoRMD formulates RNA modification prediction as a unified multi-class problem based on experimentally verified modification sites. This design not only mitigates label noise introduced by negative sample generation but also enables the model to learn discriminative representations among multiple modification types directly from real data. In addition, EvoRMD integrates data from multiple species and incorporates biological context embeddings—including organ/tissue, cell type, and subcellular localization—thereby providing a more comprehensive and biologically grounded representation of RNA modification landscapes. Such integration improves the model’s stability, generalization across species, and scalability for downstream biological applications.

To facilitate model training and evaluation, the dataset is randomly divided into training, validation and testing sets in the ratio of 8:1:1. Specifically, 0.8 of the data is used to train the model, 0.1 is used for hyperparameter tuning and performance monitoring during training, and the remaining 0.1 is used for model evaluation. This allocation ensures that the model is evaluated on unseen data while maintaining a sufficient number of samples for effective training.

#### 5.5.2 Evaluation metrics

Due to the highly unbalanced class distribution in the dataset, we used a comprehensive set of standard evaluation metrics to assess the effectiveness of the proposed method. These include accuracy (ACC), precision (Pre), sensitivity or recall (Sn), specificity (Sp), F1 score (F1), Matthews correlation coefficient (MCC), Area Under the Receiver Operating Characteristic Curve (AUC), and Area Under the Precision-Recall Curve (AUPRC). The formulas used to calculate these metrics are as follows:

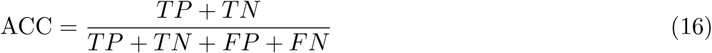

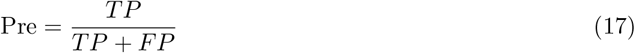

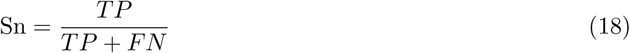

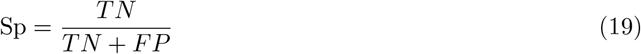

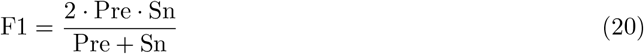

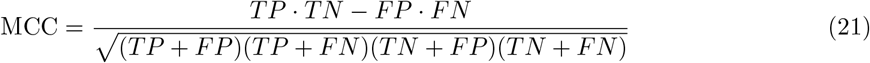

Here, True Positives (TP), True Negatives (TN), False Positives (FP), and False Negatives (FN) represent the number of correctly predicted modification types, correctly predicted non-target types, incorrectly predicted target types, and incorrectly predicted non-target types, respectively. AUC is a threshold independent metric that provides a comprehensive assessment of a model’s ability to discriminate between classes. In contrast, metrics such as Precision, Recall, and F1 require the conversion of predicted probabilities into binary predictions, which depend on a classification threshold. In our case, the optimal threshold for each class was determined by maximising its MCC score.

Furthermore, due to the highly imbalanced nature of the dataset, we adopted the area under the Precision-Recall curve (AUPRC) as a key criterion for hyperparameter selection. Compared to other metrics, AUPRC is more sensitive to performance on minority classes and provides a more informative analysis under class imbalance, making it particularly suitable for capturing the effectiveness of the model on low-frequency RNA modifications.

#### 5.5.3 Interpretation protocol

In order to analyse the decision-making mechanism and biological connotation of RNA modification prediction models, this study has developed a systematic and interpretable analysis method.

First, based on the attention mechanism, we calculate the attention weight of each nucleotide site to quantify its importance in predicting specific RNA modification categories. Specifically, for the sequences correctly predicted by the model, a sliding window of fixed length (10 nt) is used to extract subsequences and the average of the attention weights within the window is calculated. The top 10 subsequence windows with the highest attention weights were selected as candidate RNA modification conserved motifs. Furthermore, the predicted conserved motif features in each category were visualised by calculating the position weight matrix (PWM) and plotting the sequence signatures.

Second, to verify the relationship between the motifs obtained from the Attention analysis and the known RNA modification motifs, we exported the model-predicted PWMs to a MEME format file and used the Tomtom tool ^54^ in the MEME suite to compare the similarity with the known motifs in the RMBase database ^18^. The correspondence between the potential functional motifs captured by the model and the known biological motifs was clarified by significance threshold screening (p-value *<* 0.05).

Finally, to explore the potential sequence and feature commonalities among different RNA modification classes, we implemented a cross-category similarity analysis: (1) cross-comparison of known motifs across categories using the Tomtom tool ^54^; and (2) computation of Pearson correlation between attention weight matrices of different modification classes to quantify the commonality of the model’s feature capture among different RNA modification types.

The above method systematically elucidates the biological basis of the model prediction and effectively improves the credibility and biological interpretability of the prediction results.

#### 5.5.4 Computing environment

All deep learning experiments were implemented in Python using the PyTorch framework and run on a Linux workstation equipped with two NVIDIA GeForce RTX 3090 GPUs (24 GB memory per GPU), with NVIDIA driver version 535.183.01 and CUDA 12.2. Batch sizes and sequence truncation lengths were chosen such that all models fit comfortably within the available GPU memory.

## 6 Declarations

### 6.2 Ethics approval and consent to participate

Not applicable.

### 6.3 Consent for publication

Not applicable.

### 6.4 Availability of data and materials

The datasets supporting the conclusions of this article are available from the RMBase ^18^ database at https://rna.sysu.edu.cn/rmbase3/.

### 6.6 Author contributions

B.W.: Conceptualization, Investigation, Methodology, Data Curation, Software, Resources, Formal Analysis, Visualization, Validation, Writing Manuscript; H.Z.: Methodology, Resources, Manuscript Proofreading; T.C.: Methodology, Manuscript Proofreading; X.W.: Methodology, Manuscript Proof-reading; J.S. & H.X.: Conceptualization, Investigation, Methodology, Software, Resources, Supervision, Project Administration, Revising Manuscript. All authors read and approved the final manuscript.

### 6.7 Code availability

The deep learning framework was implemented using Pytorch, and the Python codes can be freely accessed at https://github.com/Gardeina/EvoRMD.

## References

[1] Cohn, W.E.: Pseudouridine, a carbon-carbon linked ribonucleoside in ribonucleic acids: Isolation, structure, and chemical characteristics. Journal of Biological Chemistry 235(5), 1488–1498 (1960) 10.1016/s0021-9258(18)69432-3

[2] Machnicka, M.A., Milanowska, K., Osman Oglou, O., Purta, E., Kurkowska, M., Olchowik, A., Januszewski, W., Kalinowski, S., Dunin-Horkawicz, S., Rother, K.M., Helm, M., Bujnicki, J.M., Grosjean, H.: Modomics: a database of rna modification pathways—2013 update. Nucleic Acids Research 41(D1), 262–267 (2012) 10.1093/nar/gks1007

[3] Sun, X.H., Zhao, L.P., Zou, Q., Wang, Z.B.: Identification of microrna genes and their mrna targets in festuca arundinacea. Applied Biochemistry and Biotechnology 172(8), 3875–3887 (2014) 10.1007/s12010-014-0805-6

[4] Hu, J., Cai, J., Xu, T., Kang, H.: Epitranscriptomic ¡scp¿mrna¡/scp¿ modifications governing plant stress responses: underlying mechanism and potential application. Plant Biotechnology Journal 20(12), 2245–2257 (2022) 10.1111/pbi.13913

[5] Wilkinson, E., Cui, Y.-H., He, Y.-Y.: Context-dependent roles of rna modifications in stress responses and diseases. International Journal of Molecular Sciences 22(4), 1949 (2021) 10.3390/ijms22041949

[6] Vu, L.P., Pickering, B.F., Cheng, Y., Zaccara, S., Nguyen, D., Minuesa, G., Chou, T., Chow, A., Saletore, Y., MacKay, M., Schulman, J., Famulare, C., Patel, M., Klimek, V.M., Garrett-Bakelman, F.E., Melnick, A., Carroll, M., Mason, C.E., Jaffrey, S.R., Kharas, M.G.: The n6-methyladenosine (m6a)-forming enzyme mettl3 controls myeloid differentiation of normal hematopoietic and leukemia cells. Nature Medicine 23(11), 1369–1376 (2017) 10.1038/nm.4416

[7] Choe, J., Lin, S., Zhang, W., Liu, Q., Wang, L., Ramirez-Moya, J., Du, P., Kim, W., Tang, S., Sliz, P., Santisteban, P., George, R.E., Richards, W.G., Wong, K.-K., Locker, N., Slack, F.J., Gregory, R.I.: mrna circularization by mettl3-eif3h enhances translation and promotes oncogenesis. Nature 561(7724), 556–560 (2018) 10.1038/s41586-018-0538-8

[8] Jin, G., Xu, M., Zou, M., Duan, S.: The processing, gene regulation, biological functions, and clinical relevance of n4-acetylcytidine on rna: A systematic review. Molecular Therapy - Nucleic Acids 20, 13–24 (2020) 10.1016/j.omtn.2020.01.037

[9] Sharma, S., Langhendries, J.-L., Watzinger, P., Kotter, P., Entian, K.-D., Lafontaine, D.L.J.: Yeast kre33 and human nat10 are conserved 18s rrna cytosine acetyltransferases that modify trnas assisted by the adaptor tan1/thumpd1. Nucleic Acids Research 43(4), 2242–2258 (2015) 10.1093/nar/gkv075

[10] Zhao, B.S., Roundtree, I.A., He, C.: Post-transcriptional gene regulation by mrna modifications. Nature reviews Molecular cell biology 18(1), 31–42 (2017)

[11] Sun, H., Li, K., Liu, C., Yi, C.: Regulation and functions of non-m6a mrna modifications. Nature Reviews Molecular Cell Biology 24(10), 714–731 (2023)

[12] Zhou, Y., Zeng, P., Li, Y.-H., Zhang, Z., Cui, Q.: Sramp: prediction of mammalian n6-methyladenosine (m6a) sites based on sequence-derived features. Nucleic Acids Research 44(10), 91–91 (2016) 10.1093/nar/gkw104 https://academic.oup.com/nar/article-pdf/44/10/e91/19694608/gkw104.pdf

[13] Fang, T., Zhang, Z., Sun, R., Zhu, L., He, J., Huang, B., Xiong, Y., Zhu, X.: Rnam5cpred: Prediction of rna 5-methylcytosine sites based on three different kinds of nucleotide composition. Molecular Therapy - Nucleic Acids 18, 739–747 (2019) 10.1016/j.omtn.2019.10.008

[14] Jiang, J., Wang, Z., Liu, Y., Wang, Z., Wang, Y., Yao, Y., Huang, Y.: m6ampred: Identifying rna n6, 2’-o-dimethyladenosine (m6am) sites based on sequence-derived information. Methods 203, 328–334 (2022) 10.1016/j.ymeth.2021.01.007

[15] Zhang, Y., Li, Y., Wang, S., Zhang, Y.: Deepbindgcn: Integrating deep learning with graph convolutional networks for protein-ligand binding affinity prediction. Briefings in Bioinformatics 21(6), 1662–1673 (2020) 10.1093/bib/bbz112

[16] Zou, Q., Xing, P., Wei, L., Liu, B.: Gene2vec: gene subsequence embedding for prediction of mammalian n6-methyladenosine sites from mrna. RNA 25(2), 205–218 (2019) 10.1261/rna.069112.118

[17] Wang, C., Ju, Y., Zou, Q., Lin, C.: Deepac4c: a convolutional neural network model with hybrid features composed of physicochemical patterns and distributed representation information for identification of n4-acetylcytidine in mrna. Bioinformatics 38(1), 52–57 (2021) 10.1093/bioinformatics/btab611

[18] Xuan, J., Chen, L., Chen, Z., Pang, J., Huang, J., Lin, J., Zheng, L., Li, B., Qu, L., Yang, J.: Rmbase v3.0: decode the landscape, mechanisms and functions of rna modifications. Nucleic Acids Research 52(D1), 273–284 (2023) 10.1093/nar/gkad1070 https://academic.oup.com/nar/article-pdf/52/D1/D273/55040098/gkad1070.pdf

[19] Boccaletto, P., Stefaniak, F., Ray, A., Cappannini, A., Mukherjee, S., Purta, E., Kurkowska, M., Shirvanizadeh, N., Destefanis, E., Groza, P., et al.: Modomics: a database of rna modification pathways. 2021 update. Nucleic acids research 50(D1), 231–235 (2022)

[20] Squires, J.E., Patel, H.R., Nousch, M., Sibbritt, T., Humphreys, D.T., Parker, B.J., Suter, C.M., Preiss, T.: Widespread occurrence of 5-methylcytosine in human coding and non-coding rna. Nucleic acids research 40(11), 5023–5033 (2012)

[21] Schwartz, S., Bernstein, D.A., Mumbach, M.R., Jovanovic, M., Herbst, R.H., Leon-Ricardo, B.X., Engreitz, J.M., Guttman, M., Satija, R., Lander, E.S., et al.: Transcriptome-wide mapping reveals widespread dynamic-regulated pseudouridylation of ncrna and mrna. Cell 159(1), 148–162 (2014)

[22] Liang, Z., Ye, H., Ma, J., Wei, Z., Wang, Y., Zhang, Y., Huang, D., Song, B., Meng, J., Rigden, D.J., et al.: m6a-atlas v2. 0: updated resources for unraveling the n 6-methyladenosine (m6a) epitranscriptome among multiple species. Nucleic acids research 52(D1), 194–202 (2024)

[23] Liu, J., Li, K., Cai, J., Zhang, M., Zhang, X., Xiong, X., Meng, H., Xu, X., Huang, Z., Peng, J., et al.: Landscape and regulation of m6a and m6am methylome across human and mouse tissues. Molecular cell 77(2), 426–440 (2020)

[24] Song, B., Huang, D., Zhang, Y., Wei, Z., Su, J., Magalhaes, J., Rigden, D.J., Meng, J., Chen, K.: m6a-tshub: unveiling the context-specific m 6 a methylation and m 6 a-affecting mutations in 23 human tissues. Genomics, proteomics & bioinformatics 21(4), 678–694 (2023)

[25] Xiong, X., Hou, L., Park, Y.P., Molinie, B., 2, G.C.A.K.G..A.F., Gregory, R.I., Kellis, M.: Genetic drivers of m6a methylation in human brain, lung, heart and muscle. Nature genetics 53(8), 1156–1165 (2021)

[26] Zhang, Y., Wang, Z., Zhang, Y., Li, S., Guo, Y., Song, J., Yu, D.-J.: Interpretable prediction models for widespread m6a rna modification across cell lines and tissues. Bioinformatics 39(12), 709 (2023)

[27] Li, X., Xiong, X., Wang, K., Wang, L., Shu, X., Ma, S., Yi, C.: Transcriptome-wide mapping reveals reversible and dynamic n 1-methyladenosine methylome. Nature chemical biology 12(5), 311–316 (2016)

[28] Ju, C.-W., Li, H., Jiang, B., Zhu, X., Cui, L., Han, Z., Zou, J., Liu, Y., Shen, S., Shah, H., et al.: Quantitative craci reveals transcriptome-wide distribution of rna dihydrouridine at base resolution. Nature Communications 16(1), 8863 (2025)

[29] Feltz, C., DeHaven, A.C., Hoskins, A.A.: Stress-induced pseudouridylation alters the structural equilibrium of yeast u2 snrna stem ii. Journal of molecular biology 430(4), 524–536 (2018)

[30] Iqbal, M.S., Abbasi, R., Bin Heyat, M.B., Akhtar, F., Abdelgeliel, A.S., Albogami, S., Fayad, E., Iqbal, M.A.: Recognition of mrna n4 acetylcytidine (ac4c) by using non-deep vs. deep learning. Applied Sciences 12(3), 1344 (2022) 10.3390/app12031344

[31] Ao, C., Zou, Q., Yu, L.: Nmrf: identification of multispecies rna 2’-o-methylation modification sites from rna sequences. Briefings in Bioinformatics 23(1) (2021) 10.1093/bib/bbab480

[32] Zhao, W., Zhou, Y., Cui, Q., Zhou, Y.: Paces: prediction of n4-acetylcytidine (ac4c) modification sites in mrna. Scientific Reports 9(1) (2019) 10.1038/s41598-019-47594-7

[33] Alam, W., Tayara, H., Chong, K.T.: Xg-ac4c: identification of n4-acetylcytidine (ac4c) in mrna using extreme gradient boosting with electron-ion interaction pseudopotentials. Scientific Reports 10(1) (2020) 10.1038/s41598-020-77824-2

[34] Chen, J., Hu, Z., Sun, S., Tan, Q., Wang, Y., Yu, Q., Zong, L., Hong, L., Xiao, J., Shen, T., King, I., Li, Y.: Interpretable rna foundation model from unannotated data for highly accurate rna structure and function predictions (2022) 10.1101/2022.08.06.503062

[35] Schaefer, M., Pollex, T., Hanna, K., Lyko, F.: Rna cytosine methylation analysis by bisulfite sequencing. Nucleic Acids Research 37(2), 12 (2009) 10.1093/nar/gkn954

[36] Song, Z., Huang, D., Song, B., Wang, Y., Zhao, F., Liu, Y., Ma, J., Xiao, Y., Zhang, Z., Yang, Y.: Attention-based multi-label neural networks for integrated prediction and interpretation of twelve widely occurring rna modifications. Nature Communications 12(1), 4011 (2021) 10.1038/s41467-021-24313-3

[37] Chen, T., Wu, T., Pan, D., Xie, J., Zhi, J., Wang, X., Quan, L., Lyu, Q.: Transrnam: Identifying twelve types of rna modifications by an interpretable multi-label deep learning model based on transformer. IEEE/ACM Transactions on Computational Biology and Bioinformatics 20(6), 36233634 (2023) 10.1109/TCBB.2023.3307419

[38] Liu, Q., Zeng, M., Li, Y., Lu, C., Kan, S., Guo, F., Li, M.: 2ome-lm: predicting 2’-o-methylation sites in human rna using a pre-trained rna language model. Bioinformatics 41(8), 417 (2025) 10.1093/bioinformatics/btaf417

[39] Suleman, M.T., Alturise, F., Alkhalifah, T., Khan, Y.D.: m1a-ensem: accurate identification of 1-methyladenosine sites through ensemble models. BioData Mining 17(1), 4 (2024) 10.1186/s13040-023-00353-x

[40] Bilal, A., Alarfaj, F.K., Khan, R.A., Suleman, M.T., Long, H.: m5c-iensem: 5-methylcytosine sites identification through ensemble models. Bioinformatics 41(1), 722 (2025) 10.1093/bioinformatics/btae722

[41] Harun-Or-Roshid, M., Maeda, K., Phan, L.T., Manavalan, B., Kurata, H.: Stack-dhupred: Advancing the accuracy of dihydrouridine modification sites detection via stacking approach. Computers in Biology and Medicine 169, 107848 (2024) 10.1016/j.compbiomed.2023.107848

[42] Gu, W.-J., Huang, Z., Chen, Z., Chen, Z., Chen, N.-T., Wei, L., Wei, Z., Jiang, X.: m5u-hybridnet: Integrating an rna foundation model with cnn features for accurate prediction of 5-methyluridine modification sites. Journal of Chemical Information and Modeling 65(14), 8079–8096 (2025) 10.1021/acs.jcim.4c01847

[43] Song, B., Shi, G., Zhang, W., Zhang, L., Yang, Y., Chen, K., Zhou, Y., Zhang, Q., Huang, D., Liu, G., Yang, Y., Zhang, W., Wang, J., Zhou, Y.: m6a-tshub: Unveiling the context-specific m6a methylation and m6a-affecting mutations in 23 human tissues. Genomics, Proteomics & Bioinformatics 21(4), 678–695 (2023) 10.1016/j.gpb.2022.09.001

[44] Yu, J., Gao, W., Chen, S., Lu, R., Qiao, J., Jin, J., Wei, L., Shi, H., Zhang, Z., Cui, F., Jiang, X., Yan, Z.: Mcamef-bert: an efficient deep learning method for rna n7-methylguanosine site prediction via multi-branch feature integration. Briefings in Bioinformatics 26(5), 447 (2025) 10.1093/bib/bbaf447

[45] Han, B., Bai, S., Liu, Y., Wu, J., Feng, X., Xin, R.: Definer: A computational method for accurate identification of rna pseudouridine sites based on deep learning. PLOS ONE 20(4), 0320077 (2025) 10.1371/journal.pone.0320077

[46] Cui, L., Ma, R., Cai, J., Guo, C., Chen, Z., Yao, L., Wang, Y., Fan, R., Wang, X., Shi, Y.: Rna modifications: importance in immune cell biology and related diseases. Signal Transduction and Targeted Therapy 7(1), 334 (2022) 10.1038/s41392-022-01175-9

[47] Liu, J., Yue, Y., Han, D., Wang, X., Fu, Y., Zhang, L., Jia, G., Yu, M., Lu, Z., Deng, X., Dai, Q., Chen, W., He, C.: Biological roles of adenine methylation in rna. Nature Reviews Genetics 24(2), 143–160 (2023) 10.1038/s41576-022-00534-0

[48] Sloan, K.E., Warda, A.S., Sharma, S., Entian, K.-D., Lafontaine, D.L.J., Bohnsack, M.T.: Tuning the ribosome: The influence of rrna modification on eukaryotic ribosome biogenesis and function. RNA Biology 14(9), 1138–1152 (2017) 10.1080/15476286.2016.1259781

[49] Anderson, J., Phan, L., Cuesta, R., Carlson, B.A., Pak, M., Asano, K., Bjork, G.R., Tamame, M., Hinnebusch, A.G.: Genome-wide analysis of n1-methyl-adenosine modification in yeast trna. RNA 24(3), 361–373 (2018) 10.1261/rna.2057810

[50] Malbec, L., Zhang, T., Chen, Y.S., Zhang, Y., Sun, B.F., Shi, B.Y., Zhao, Y.L., Yang, Y., Wang, X., Jin, H., Wang, X., Wang, Y., Wang, X., Zhang, X., Zhou, J., Yang, Y.G.: Dynamic methylome of internal mrna n7-methylguanosine and its regulatory role in translation. Cell Research 29(11), 927–941 (2019) 10.1038/s41422-019-0230-z

[51] Karijolich, J., Yu, Y.-T.: Converting nonsense codons into sense codons by targeted pseudouridylation. Nature 474(7351), 395–398 (2011) 10.1038/nature10165

[52] Jia, R., Xie, Y., Wen, J., Guo, J., Wang, X., Wang, X., Zhang, Z., Shi, H.: Comprehensive functional annotation of mrna methylation reveals context-dependent roles in gene regulation and cancer. eLife 13, 92537 (2024) 10.7554/eLife.92537.3

[53] Linder, B., Grozhik, A.V., Olarerin-George, A.O., Meydan, C., Mason, C.E., Jaffrey, S.R.: Single-nucleotide-resolution mapping of m6a and m6am throughout the transcriptome. Nature Methods 12(8), 767–772 (2015) 10.1038/nmeth.3453

[54] Gupta, S., Stamatoyannopoulos, J.A., Bailey, T.L., Noble, W.S.: Quantifying similarity between motifs. Genome Biology 8(2), 24 (2007) 10.1186/gb-2007-8-2-r24

[55] Song, P., Yi, C.: Interplays of different types of epitranscriptomic mrna modifications. Trends in Genetics 37(11), 1028–1045 (2021) 10.1016/j.tig.2021.09.003

[56] Li, X., Zhang, Y., Wang, B., Chen, Y., Liu, W., Xu, Z., Wang, Y., Yang, Y.: Transcriptome-wide mapping reveals dynamic co-modification of mrna by multiple rna modifications. Nature Communications 14(1), 2013 (2023) 10.1038/s41467-023-37670-y

[57] Mauer, J., Luo, X., Blanjoie, A., Jiao, X., Grozhik, A.V., Patil, D.P., Linder, B., Pickering, B.F., Vasseur, J.-J., Chen, Q., Jaffrey, S.R.: Reversible methylation of m6am in the 5’ cap controls mrna stability. Nature 541(7637), 371–375 (2017) 10.1038/nature21022

[58] Li, Q., Zhu, Q.: The role of demethylase alkb homologs in cancer. Frontiers in Oncology 13, 1153463 (2023) 10.3389/fonc.2023.1153463

[59] Zhang, X., Liu, H., Lin, Y., Li, Q., Wang, M., Zhang, W., Chen, Q., Xu, J., Zhao, Y., Liu, M., Yang, G.: Targeting ybx1-m5c mediates rnf115 mrna circularisation and translation to enhance vulnerability of ferroptosis in hepatocellular carcinoma. Cell Reports 42(7), 113127 (2024) 10.1016/j.celrep.2024.113127

[60] Arzumanian, V.A., Timofeeva, E.V., Kiseleva, O.I., Poverennaya, E.V.: Drug-metabolizing gene expression identity: Comparison across liver tissues and model cell lines. Biomedicines 13(11) (2025)

[61] Huggett, Z.J., Smith, A., De Vivo, N., Gomez, D., Jethwa, P., Brameld, J.M., Bennett, A., Salter, M.: A comparison of primary human hepatocytes and hepatoma cell lines to model the effects of fatty acids, fructose and glucose on liver cell lipid accumulation. Nutrients 15(1) (2023)

[62] Wisniewska, M.J., Wencel, A., Jakubowska, M., Motyl, J.A., Dudek, K., Burzynska, B., Pijanowska, D.G., Pluta, K.D.: Variable expression of hepatic genes in different liver tumor cell lines: conclusions for drug testing. Frontiers in Cell and Developmental Biology Volume 13 - 2025 (2025) 10.3389/fcell.2025.1646602

[63] Arzumanian, V., Pyatnitskiy, M., Poverennaya, E.: Comparative transcriptomic analysis of three common liver cell lines. International Journal of Molecular Sciences 24(10), 8791 (2023) 10.3390/ijms24108791

[64] Gunn, P.J., Green, C.J., Pramfalk, C., Hodson, L.: In vitro cellular models of human hepatic fatty acid metabolism: differences between huh7 and hepg2 cell lines in human and fetal bovine culturing serum. Physiological Reports 5(24), 13532 (2017) 10.14814/phy2.13532

[65] Zhang, C., Yang, T., Chen, H., Ding, X., Chen, H., Liang, Z., Zhao, Y., Ma, S., Liu, X.: Mettl3 inhibition promotes radiosensitivity in hepatocellular carcinoma through regulation of slc7a11 expression. Cell Death & Disease 16(1), 9 (2025) 10.1038/s41419-024-07317-x

[66] Wang, F., Hu, Y., Wang, H., et al.: Lncrna fto-it1 promotes glycolysis and progression of hepatocellular carcinoma through modulating fto-mediated n6-methyladenosine modification on glut1 and pkm2. Journal of Experimental & Clinical Cancer Research 42(1), 267 (2023) 10.1186/s13046-023-02847-2

[67] Li, J., Zhu, L., Shi, Y., Liu, J., Lin, L., Chen, X.: m6a demethylase fto promotes hepatocellular carcinoma tumorigenesis via mediating pkm2 demethylation. American Journal of Translational Research 11(9), 6084–6092 (2019)

[68] Dai, Y.Z., Liu, Y.D., Li, J., Chen, M.T., Huang, M., Wang, F., Yang, Q.S., Yuan, J.H., Sun, S.H.: Mettl16 promotes hepatocellular carcinoma progression through downregulating rab11b-as1 in an m6a-dependent manner. Cellular and Molecular Biology Letters 27(1), 41 (2022) 10.1186/s11658-022-00342-8

[69] Huang, W., Chen, T.Q., Fang, K., et al.: N6-methyladenosine methyltransferases: functions, regulation, and clinical potential. Journal of Hematology & Oncology 14(1), 117 (2021) 10.1186/s13045-021-01129-8

[70] Safra, M., Sas-Chen, A., Nir, R., Farouq, D., Vasseur, J.-J., Debart, F., Levanon, E.Y., Schwartz, S.: The m1a landscape on cytosolic and mitochondrial mrna at single-nucleotide resolution. Nature 551(7679), 251–255 (2017) 10.1038/nature24456

[71] Li, X., Xiong, X., Wang, K., Wang, L., Shu, X., Ma, S., Yi, C.: Transcriptome-wide mapping reveals reversible and dynamic n1-methyladenosine methylome in mammalian mrna. Cell Research 27(5), 616–624 (2017) 10.1038/cr.2017.55

[72] Dominissini, D., Moshitch-Moshkovitz, S., Schwartz, S., Salmon-Divon, M., Ungar, L., Osenberg, S., Cesarkas, K., Jacob-Hirsch, J., Amariglio, N., Kupiec, M., Sorek, R., Rechavi, G.: Topology of the human and mouse m6a rna methylomes revealed by m6a-seq. Nature 485, 201–206 (2012) 10.1038/nature11112

[73] Gillespie, M., Jassal, B., Stephan, R., Milacic, M., Rothfels, K., Senff-Ribeiro, A., Griss, J., Sevilla, C., Matthews, L., Gong, C., Deng, C., Varusai, T., Ragueneau, E., Haider, Y., May,, Shamovsky, V., Weiser, J., Brunson, T., Sanati, N., Beckman, L., Shao, X., Fabregat, A., Sidiropoulos, K., Murillo, J., Viteri, G., Cook, J., Shorser, S., Bader, G., Demir, E., Sander, C., Haw, R., Wu, G., Stein, L., Hermjakob, H., D’Eustachio, P.: The reactome pathway knowledgebase 2022. Nucleic Acids Research 50(D1), 687–692 (2021) 10.1093/nar/gkab1028 https://academic.oup.com/nar/article-pdf/50/D1/D687/42058295/gkab1028.pdf

[74] Bressac, B., Galvin, K.M., Liang, T.J., Isselbacher, K.J., Wands, J.R., Ozturk, M.: Abnormal structure and expression of p53 gene in human hepatocellular carcinoma. Proceedings of the National Academy of Sciences 87(5), 1973–1977 (1990) 10.1073/pnas.87.5.1973

[75] Hsu, I.C., Metcalf, R.A., Sun, T., Welsh, J.A., Wang, N.J., Harris, C.C.: Mutational hotspot in the p53 gene in human hepatocellular carcinomas. International Journal of Cancer 55(3), 397–401 (1993) 10.1002/ijc.2910550310

[76] Guo, L., Dial, S., Shi, L., Branham, W., Liu, J., Fang, J.-L., Green, B., Deng, H., Kaput, J., Ning, B.: Similarities and differences in the expression of drug-metabolizing enzymes between human hepatic cell lines and primary human hepatocytes. Drug Metabolism and Disposition 39(3), 528–538 (2011) 10.1124/dmd.110.035873

[77] Buszczak, M., Signer, R.A.J., Morrison, S.J.: Protein synthesis and the control of stem cell fate. Cell 159(2), 242–251 (2014) 10.1016/j.cell.2014.09.024

[78] Tieh, T.H., Zhao, Y., Shah, N.M., Heng, H.L., Tom, E., Yeo, A., Zhao, Y., Zeng, Q., So, J.B., Wan, W.K.: Constitutive activation of FGFR4 signaling supports the growth and survival of glioblastoma multiforme cells. Oncotarget 5(7), 1794–1806 (2014) 10.18632/oncotarget.1752

[79] Venkatesh, H.S., Johung, T.-B., Caretti, V., Noll, A., Tang, Y., Nagaraja, S., Gibson, E.M., Mount, W., Polepalli, J., Mitra, S.S., et al.: Neuronal activity promotes glioma growth through neuroligin-3 secretion. Cell 161(4), 803–816 (2015) 10.1016/j.cell.2015.04.012

[80] Fu, L., Niu, B., Zhu, Z., Wu, S., Li, W.: Cd-hit: accelerated for clustering the next-generation sequencing data. Bioinformatics 28(23), 3150–3152 (2012) 10.1093/bioinformatics/bts565

[81] Li, W., Godzik, A.: Cd-hit: a fast program for clustering and comparing large sets of protein or nucleotide sequences. Bioinformatics 22(13), 1658–1659 (2006) 10.1093/bioinformatics/btl158

