## Appendix A for "EvoRMD: Integrating Biological Context and Evolutionary RNA Language Models for Interpretable Prediction of RNA Modifications": Appendix.pdf

#### A.1 Position Weights with Significant Influence on Model Predictions

**Table A1**: **Top 8 attention positions and average N21 attention for each RNA modification.** The table summarizes the attention weights extracted from the attention module, highlighting the top eight nucleotide positions with the highest weights for each modification type. Additionally, the attention weight corresponding to position 21—the designated modification site—is specifically reported.

| Type | Top 8 Positions | Attention Weights | N21 Attention |
|---|---|---|---|
| Am | [20, 22, 4, 26, 29, 24, 14, 6] | [0.0304, 0.0273, 0.0268, 0.0267, 0.0266, 0.0260, 0.0257, 0.0256] | 0.0221 |
| Cm | [15, 19, 3, 37, 4, 16, 6, 17] | [0.0283, 0.0274, 0.0267, 0.0265, 0.0265, 0.0264, 0.0261, 0.0257] | 0.0232 |
| D | [20, 26, 4, 19, 6, 29, 25, 27] | [0.0385, 0.0356, 0.0331, 0.0316, 0.0315, 0.0304, 0.0303, 0.0302] | 0.0113 |
| Gm | [7, 20, 26, 4, 35, 37, 6, 10] | [0.0277, 0.0277, 0.0264, 0.0260, 0.0252, 0.0252, 0.0250, 0.0247] | 0.0234 |
| $m^1A$ | [20, 22, 33, 14, 28, 6, 4, 2] | [0.0270, 0.0265, 0.0263, 0.0261, 0.0258, 0.0257, 0.0257, 0.0257] | 0.0238 |
| $m^5C$ | [7, 15, 3, 4, 16, 37, 6, 9] | [0.0295, 0.0264, 0.0263, 0.0260, 0.0259, 0.0259, 0.0256, 0.0254] | 0.0235 |
| $m^5U$ | [20, 24, 19, 12, 13, 16, 41, 6] | [0.0465, 0.0394, 0.0371, 0.0310, 0.0308, 0.0307, 0.0270, 0.0266] | 0.0122 |
| $m^6A$ | [18, 20, 23, 19, 30, 25, 26, 17] | [0.0325, 0.0283, 0.0270, 0.0265, 0.0258, 0.0257, 0.0256, 0.0255] | 0.0213 |
| $m^7G$ | [7, 27, 20, 15, 26, 37, 4, 29] | [0.0283, 0.0268, 0.0265, 0.0264, 0.0256, 0.0246, 0.0245, 0.0244] | 0.0228 |
| Um | [7, 15, 29, 25, 20, 6, 27, 34] | [0.0349, 0.0303, 0.0283, 0.0282, 0.0262, 0.0260, 0.0256, 0.0253] | 0.0141 |
| Y | [7, 27, 15, 29, 6, 25, 3, 5] | [0.0353, 0.0299, 0.0285, 0.0279, 0.0274, 0.0261, 0.0260, 0.0256] | 0.0176 |

### Appendix B

The following supplementary files are provided with this manuscript:

- **Appendix File A: All Tomtom alignment results of the conserved motifs discovered by EvoRMD and those in the RMBase database.**
- **Appendix File B: Similarity comparison of conserved motifs with different modifications in EvoRMD.**
- **Appendix File C: Similarity comparison of conserved motifs with different modifications in RMBase.**

### Appendix C Evaluation of Ensemble Learning Strategy for Handling Class Imbalance

To mitigate potential information loss from downsampling, we evaluated a Bagging-based ensemble of five EvoRMD models trained on distinct subsets of majority classes. Unexpectedly, the ensemble approach degraded performance, with the Overall MCC dropping from 97.82% (single model, $\alpha = 0.6$) to 85.93% (Table A2). We attribute this decline to the divergent evolution of feature spaces during RNA-FM fine-tuning, where averaging logits from biologically distinct latent representations caused "destructive interference," blurring decision boundaries for subtle classes like $m^1$A. Additionally, the high motif redundancy in abundant classes meant that introducing extra samples provided low marginal utility while overwhelming signals from minority classes. Given the five-fold increase in computational cost without performance gains, we conclude that a single model trained on a representative dataset offers superior coherence and predictive accuracy.

**Table A2**: Performance comparison between the Original Single Model and the Bagging Ensemble ($K = 5$) approach. The table reports the Matthews Correlation Coefficient (MCC %) for each modification type. The best result in each row is highlighted in bold.

| **Modification** | **Original Single Model (MCC %)** | **Ensemble (K=5) (MCC %)** | **Performance Gap (pp)** |
|---|---|---|---|
| Am | **75.97** | 51.47 | -24.50 |
| Cm | **94.46** | 83.64 | -10.82 |
| D | **87.02** | 74.01 | -13.01 |
| Gm | 97.24 | **97.81** | +0.57 |
| $m^1A$ | **92.12** | 41.58 | -50.54 |
| $m^5C$ | **98.90** | 95.96 | -2.94 |
| $m^5U$ | 70.69 | **78.29** | +7.60 |
| $m^6A$ | **99.08** | 86.72 | -12.36 |
| $m^7G$ | **98.79** | 92.80 | -5.99 |
| Um | **99.65** | 99.04 | -0.61 |
| Y | **92.76** | 84.88 | -7.88 |
| Overall | **97.82** | 85.93 | -11.89 |

*Note:* Performance Gap is calculated as the absolute difference in percentage points (Ensemble - Original). The gray row indicates the weighted overall performance.
